# Synaptic vesicles that store monoamines and glutamate differ in protein composition

**DOI:** 10.1101/2025.05.06.651945

**Authors:** Hrach Asmerian, Alexia J. Diaz, Hongfei Xu, Juan A. Oses-Prieto, Jacob Alberts, Anna Sanetra, Barathan Gnanabharathi, Noah Carr, Alma L. Burlingame, Robert H. Edwards, Kätlin Silm

## Abstract

Neuromodulators such as the monoamines are known to differ from classical neurotransmitters like glutamate in the time scale of signaling due to activation of slower G protein-coupled receptors. Recent work has suggested that the mode of release also differs between classical and modulatory transmitters. Although many components of neurotransmitter release machinery have been identified, we still understand little about the mechanisms responsible for differences in release. In this study, we address the differences between release of dopamine and glutamate by comparing the composition of synaptic vesicles (SVs) that contain the vesicular monoamine transporter 2 (VMAT2) versus vesicular glutamate transporter 2 (VGLUT2). Previous work has shown that these SV populations differ in frequency dependence, recycling kinetics and biogenesis. Taking advantage of a CRISPR-generated knock-in mouse with a cytoplasmic hemagglutinin (HA) tag at the N-terminus of VMAT2 to immunoisolate monoamine SVs, we find differences in the abundance and isoform expression of many SV protein families. Validation in primary neurons and in brain tissue confirms these differences in SV protein abundance between dopamine and glutamate release sites. Functional analysis reveals that the loss of differentially expressed SCAMP5 selectively impairs the recycling of VGLUT2 SVs, sparing VMAT2 vesicles in the same neuronal population. These findings provide new insights into the molecular diversity of SVs and the mechanisms that regulate the release of dopamine and glutamate, with implications for the physiological role of these transmitters and behavior.

## INTRODUCTION

Considerable work over decades has elucidated the basic machinery of neurotransmitter release^1–3^ but many questions remain about the molecular mechanisms that account for the diversity in release properties at different synapses. Differences in release have been observed between neuronal populations, but also individual release sites made by the same axon, and even between vesicles in the same terminal^4–12^. This points to contribution from cell-wide differences in the expression of synaptic proteins^13–16^ but also to local mechanisms at synaptic sites, which can be further modulated by postsynaptic neurons^4, 6, 8, 10, 17^. Furthermore, most of our understanding of neurotransmitter release derives from work conducted in neurons that use fast, synaptic signaling. In contrast to classical transmitters like glutamate and GABA which activate ionotropic receptors, neuromodulators such as monoamines are often released at sites without a direct postsynaptic target^18^ and activate G-protein coupled receptors which signal over longer times scales^19^. Despite critical roles in normal brain function, the molecular regulation of the release of dopamine and other modulatory monoamine transmitters remains poorly understood.

While dopamine neurons fire tonically at low frequencies, dopamine release is significantly increased during high frequency burst firing, and this is central to reinforcement learning^20^. This property of dopamine release requires the recruitment of synaptic vesicles (SVs) specifically at high frequency. Indeed, it has been shown that dopamine release is more loosely coupled to Ca^++^ entry than fast synaptic release^21, 22^, consistent with a lower probability of release in response to isolated stimuli, and a large increase with the accumulation of presynaptic Ca^++^ at high frequency. Recent evidence also suggests that dopamine is only released from a subset of axonal varicosities^23^, which could reflect the sparsity of active zones in dopamine axons^24–26^. These findings suggest that the machinery for release in dopamine neurons differs from the standard model at classical synaptic sites.

Beyond the cell type specific features of dopamine axons, recent work in neurons that release both dopamine and glutamate has shown that the two transmitters differ in release even from the same neuron^22^. This is mediated by their storage in functionally distinct SVs that differ in synaptic recycling kinetics and in frequency dependence of fusion, which leads to faster depression of glutamate release^22^. Indeed, the formation of SVs that store monoamines relies heavily on a pathway that involves the adaptor protein complex 3 (AP-3)^22, 27^. While previous work that addressed the role of this adaptor in other neuronal populations found a minor effect on the SV cycle^28, 29^, loss of AP-3 leads to a severe reduction in the synaptic recycling of pHluorin-tagged vesicular monoamine transporter 2 (VMAT2)^22, 27^. This defect in turn reduces dopamine release in the dorsal striatum and is especially pronounced at high stimulation frequencies^27^. Indeed, these results are compatible with observations from multiple studies demonstrating that AP-3-dependent SVs preferentially fuse during high frequency activity^27, 30, 31^. Strikingly, the recycling of the vesicular glutamate transporter 2 (VGLUT2) that mediates glutamate corelease in dopamine neurons is not affected by the loss of AP-3^22, 27, 30, 32–34^. This suggests that VGLUT2 is excluded from AP3-dependent SVs and targets to a subpopulation of vesicles that likely form via the classical AP2-dependent mechanism^29, 35^. AP-3-independent vesicles also contribute to tonic dopamine release during low levels of activity as demonstrated by the more modest effect of the AP-3 knock-out on dopamine release with stimulation at low frequency^27^. Differential reliance of VMAT2 and VGLUT2 on the AP-3 pathway was also shown to extend beyond neurons that release multiple transmitters^12, 27, 30^. Compatibly with this model, VGLUT2-dependent glutamate release is consistently found to have high initial probability and to undergo depression with sustained activity^5, 7, 30, 36, 37^. The differential reliance on AP-3 suggests that differences in the biogenesis of VMAT2^+^ versus VGLUT2^+^ SVs contribute to the differences in release, likely by affecting SV composition.

Compositional diversity among SVs has previously been addressed by comparing vesicles that store the principal fast excitatory and inhibitory transmitters glutamate and GABA^38^. This comparison identified few differentially expressed proteins, suggesting a relatively uniform composition of SVs across neuronal populations. In the present study, we focus on SV populations that contain VMAT2 or VGLUT2 but also differ in biogenesis mechanism and release properties.

## RESULTS

### Proteome of VMAT2^+^ and VGLUT2^+^ synaptic vesicles

Due to the limited reliability of VMAT2 antibodies, we used CRISPR to generate a knock-in mouse with a hemagglutinin (HA) tag at the N-terminus of VMAT2 (Figure 1A)^39^. HA-VMAT2 signal detected via anti-HA antibodies is present in the cell bodies of dopamine, norepinephrine and serotonin neurons (Figure S1A, S1C-D) as well as in the axonal projections of monoamine neurons in the striatum (Figure 1A, S1B). In cultured neurons, HA-VMAT2 displays punctate signal in the processes of TH-positive neurons (Figure S1E). HA-tagged VMAT2 is therefore faithfully targeted in monoamine neurons.

**Figure 1.**
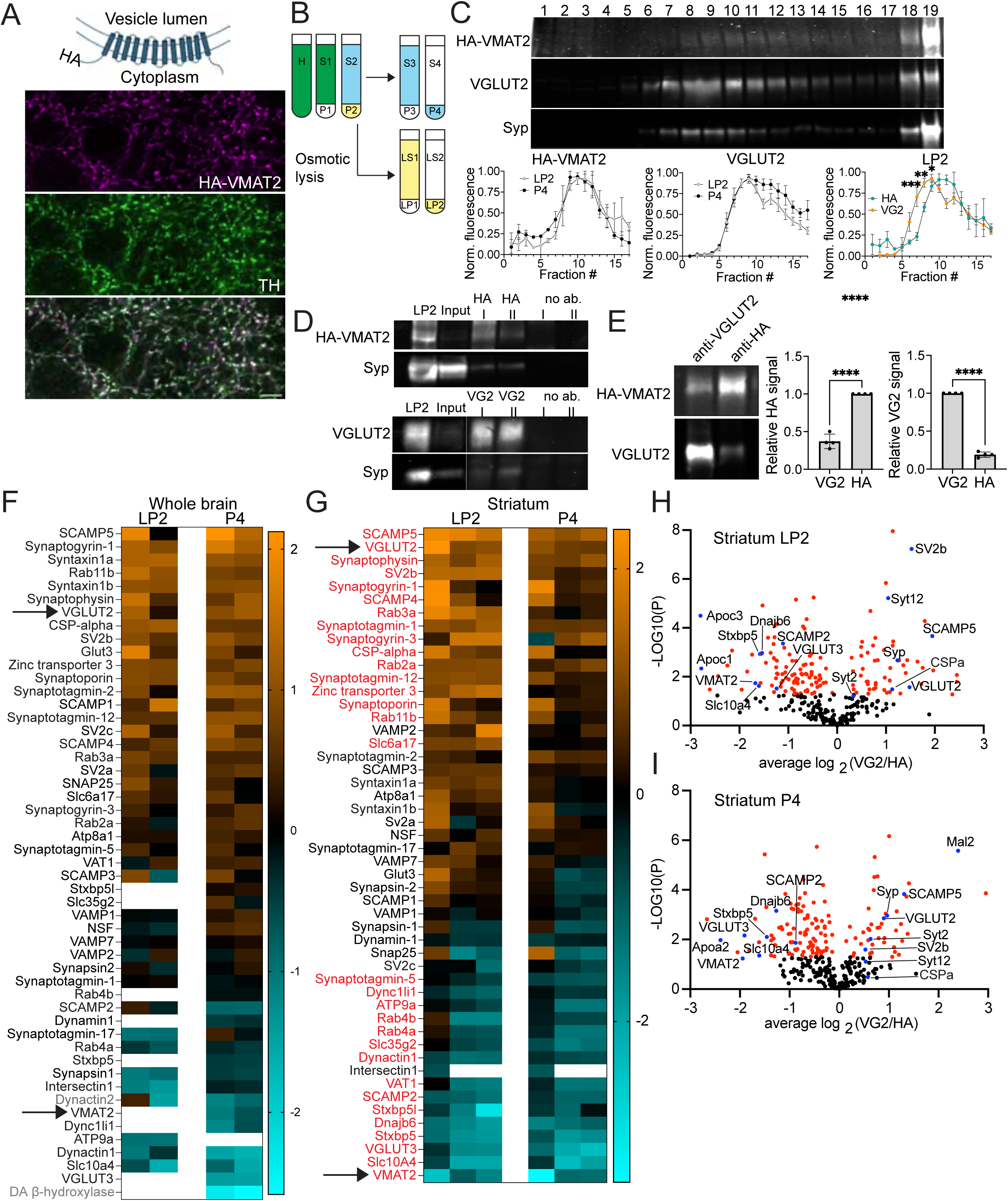
Isolation and proteomic profiling of VMAT2^+^ and VGLUT2^+^ SVs. **A.** We used CRISPR to generate a knock-in mouse with an HA tag in the cytoplasmic N-terminus of VMAT2. Confocal image of the striatum stained for HA and TH shows HA-VMAT2 signal in TH-positive axonal fibers. Scale bar = 5 µm. **B.** Scheme of the protocol for isolating synaptic vesicles. Synaptosomes from P2 pellet fraction were subjected to osmotic lysis to release synaptic vesicles from presynaptic terminals. Released vesicles (LS1) were separated from synaptic plasma membranes (LP1) and then sedimented in the LP2 pellet. In parallel, we collected light organelles from supernatant S2 to isolate axonal vesicles that were not concentrated in varicosities (P4). Fractions highlighted in color were used in the following steps. **C.** Western blot of HA-VMAT2, VGLUT2 and synaptophysin signal in LP2 samples loaded on a 5-25% glycerol gradient. 10 µL of a 500 µL fraction was loaded per lane. Graphs show the comparison of mean HA-VMAT2 and VGLUT2 signal intensities in gradient fractions of LP2 and P4 samples and the comparison of HA-VMAT2 versus VGLUT2 signal in LP2 samples quantified on dot blots. Data indicate mean ± SEM. * < 0.05, ** < 0.001, *** <0.001, by unpaired t-test **D.** Immunoblots showing HA-VMAT2 and synaptophysin (left) or VGLUT2 and synaptophysin (right) in LP2, starting material for vesicle isolation (LP2 after gradient purification and dialysis) and on Dynabeads with antibody used for isolation or in a negative control where antibody was not added. **E.** Immunoblots and quantification of HA-VMAT2 and VGLUT2 signal in vesicles immunoisolated using anti-VGLUT2 or anti-HA antibodies. Data indicate mean ± SEM. **** <0.0001, by unpaired t-test **F.** Examples of proteins quantified on VMAT2 and VGLUT2 SVs isolated from whole brain samples from HA-VMAT2 KI mice. Data was normalized to the amount of total protein in each sample and is presented as a heat map of log2 values of VGLUT2/VMAT2 ratio. Each column represents an independent experiment. Proteins indicated in gray are not detected in samples from the striatum. **G.** Examples of proteins quantified on VMAT2 and VGLUT2 SVs isolated from striatal samples of HA-VMAT2 KI mice. Data was normalized to the amount of total protein in each sample and is presented as a heat map of log2 values of VGLUT2/VMAT2 ratio. Each column represents an independent experiment. Proteins highlighted in red show significant enrichment in one SV population (P < 0.05) and have a VGLUT2/VMAT2 ratio above 1.5 or below −1.5. **H-I**. Volcano plots showing protein abundance in VMAT2 versus VGLUT2 SVs in striatal LP2 (H) and P4 samples (I) as determined by TMT labeling and mass spectrometry. Data was analyzed in three independent experiments. Data points highlighted in red have p-values < 0.05. Blue data points are associated with labels.

To isolate the vesicle populations from the HA-VMAT2 KI mice, we used a standard differential centrifugation procedure to extract synaptic vesicles from synaptosomes via osmotic lysis (Figure 1B)^30, 40^. However, since VMAT2 SVs in dopamine neurons are reported to localize in non-synaptic sites that may be excluded from synaptosomes^41^, we also collected light membranes released from axonal fragments contained in the supernatant of the P2 synaptosome pellet (Figure 1B). SVs were further purified using a 5-25% glycerol gradient where the HA-VMAT2 peak appeared in fractions corresponding to SV proteins (Figure 1C). Vesicles isolated from LP2 and P4 samples were detected in similar fractions (Figure 1C). However, we do observe a shift in HA-VMAT2 signal towards lower fractions compared to VGLUT2, which is consistent with recent reports of larger SVs in dopamine neurons^42^ (Figure 1C). VMAT2 and VGLUT2 SVs were then immunoisolated from gradient purified SVs using antibodies against VGLUT2 or the HA tag in VMAT2 (Figure 1D). Both HA-VMAT2 and VGLUT2 showed significant enrichment in SV populations isolated using their respective antibodies (Figure 1E).

To compare the protein composition of the two SV populations, we performed mass spectrometry with isotopic labeling by tandem mass tags (TMT). We initially completed this experiment in whole brain samples, which contain VMAT2 vesicles from all monoamine projections. Mass spectrometry of these samples indicates that many classical SV proteins were more abundant in the VGLUT2^+^ vesicle population while VMAT2^+^ SVs were enriched in orphan transporter Slc10A4, which is expressed by cholinergic and aminergic neurons and has a role in dopamine homeostasis^43^, and in dopamine β-hydroxylase (DBH), a luminal SV enzyme that converts dopamine into norepinephrine (Figure 1F). We also found the enrichment of VGLUT3 in monoamine vesicles, which is consistent with the presence of this glutamate transporter in serotonin neurons where it mediates glutamate corelease^44^. While the detection of DBH and Slc10A4 support the enrichment of monoamine vesicles by VMAT2 immunoisolation, many novel proteins that showed enrichment in these vesicles, as well as VMAT2 itself, were detected in only a subset of samples (Figure 1F). This presumably reflected the insufficient enrichment of this SV population from a sample where monoamine vesicles are relatively sparse. We therefore repeated the immunoisolation and subsequent analysis using striatal samples where the VMAT2^+^ SVs are more abundant due to the dense dopaminergic projection. Similarly to the whole brain, we find that many canonical SV proteins that are thought to be ubiquitously expressed in all SVs, show enrichment in VGLUT2 vesicles (summarized in Figure 1G-I). VGLUT2-enriched proteins include neuroprotective chaperone cysteine string protein α (CSPα), the synaptophysin family (synaptophysin, synaptogyrin1 and synaptoporin), as well as SCAMP4 and 5. We also find that members of the synaptotagmin (syt) protein family, which includes the fast SV calcium sensors syt1 and syt2, differ in abundance in the two SV populations. Specifically, VGLUT2+ SVs appear enriched in the fast calcium sensor syt2 and in calcium insensitive syt12 that has been implicated in a protein kinase A-dependent form of presynaptic plasticity^45–47^. The analysis of striatal samples also revealed more proteins with high abundance in VMAT2+ SVs. These include inhibitory SNARE proteins tomosyn-1 and -2 (stxbp5 and stxbp5l) that were shown to increase the barrier for SV fusion, thereby lowering release probability^48, 49^. We also find that VMAT2 vesicles contain more of chaperone DNAJb6 that is implicated in protection against α-synuclein pathology^50, 51^ and in juvenile-onset parkinsonism^52^ as well as SCAMP2 that was shown to influence trafficking of the dopamine transporter^53^. While many well-known SV proteins differ in localization to the two vesicle populations, essential components of the release machinery such as VAMP2, SNAP25 and NSF are present at comparable levels in VMAT2^+^ and VGLUT2^+^ SVs (Figure 1F, 1G). Similarly, subunits of the *v*-ATPase show no consistent enrichment in either vesicle population in striatal samples (Figure S2A).

In addition to classical SV proteins, we find the differential expression of several Rab proteins, with Rab3 and Rab11b enriched in VGLUT2^+^ SVs and Rab4 in the VMAT2^+^ sample, which could reflect trafficking of the two transporters via distinct endosomal compartments (Figure 1F, 1G). Furthermore, we observe the more abundant association of VMAT2^+^ SVs with dynactin and dynein subunits, suggesting a differential requirement for retrograde axonal and anterograde dendritic transport that would be consistent with the known release of monoamines from dendrites as well as axons (Figure 1F, 1G)^54^. Since VMAT2 and VGLUT2 vesicles differ in reliance on adaptor protein complexes for recycling from the plasma membrane, we also analyzed the data for AP-2 and AP-3 subunit abundance in the two SV populations (Figure S2B). While AP-2 subunits are present at comparable levels in both vesicle populations, AP-3 components are detected consistently only in whole brain samples where they associate preferentially with VMAT2 vesicles.

### Expression of SV proteins in primary dopamine and glutamate neurons

To assess differences in VMAT2^+^ and VGLUT2^+^ SV composition, we first used primary midbrain cultures from HA-VMAT2 KI mice as these cultures contain both dopamine neurons and VGLUT2^+^ glutamate neurons. To compare SV protein abundance in these neuronal populations, we identified regions of interest (ROIs) based on the expression of HA-VMAT2 or VGLUT2 and quantified the fluorescence intensity of candidate proteins in both sets of ROIs. Since VMAT2 localizes to the somatodendritic as well as axonal compartment^55^, we also examined the colocalization of HA-VMAT2 with the dendritic protein MAP2. We find a more diffuse HA-VMAT2 signal in MAP2^+^ fibers, while the vast majority of bright punctate structures similar in size and appearance to VGLUT2^+^ varicosities localize to MAP2^−^ axonal projections (Figure S3). These results therefore compare axonal varicosities from the two neuronal populations. As predicted by the proteomics results, orphan transporter Slc10A4^43^ localizes to VMAT2^+^ rather than VGLUT2^+^ terminals (Figure 2A). In contrast, VGLUT2^+^ sites contain more of the canonical SV protein synaptophysin (Figure 2E, S4A) as well as the related synaptogyrin1 (Figure 2F, S4B). VGLUT2^+^ terminals also contain more of presynaptic chaperone CSPα (Figure 2D) and SCAMP5 (Figure 2G, S4C). However, the SV SNARE protein VAMP2 occurs at similar levels in both release sites (Figure 2H, S4D) supporting the specificity of differences for other proteins and arguing against substantial differences in SV cluster size. For synaptophysin, synaptogyrin1, CSPα, SCAMP5 and Slc10A4, the same trends in expression were consistently observed in all analyzed dopamine and glutamate neurons, including fields of view that contain both dopamine and VGLUT2^+^ glutamate axons. However, the analysis of syt2 revealed that although syt2 signal in dopamine neurons was consistently close to background, glutamate neurons fall into two populations: VGLUT2 neurons with low or nonexistent syt2 signal similar to dopamine neurons, and another with strong syt2 expression that colocalizes with VGLUT2 (Figure 2C, 2I, 2J). These findings suggest a molecular heterogeneity among the VGLUT2^+^ glutamate neurons present in midbrain cultures. The other fast calcium sensor syt1 occurs in both types of neurons, but its signal intensity in midbrain cultures is consistently higher at dopamine release sites (Figure 2B).

**Figure 2.**
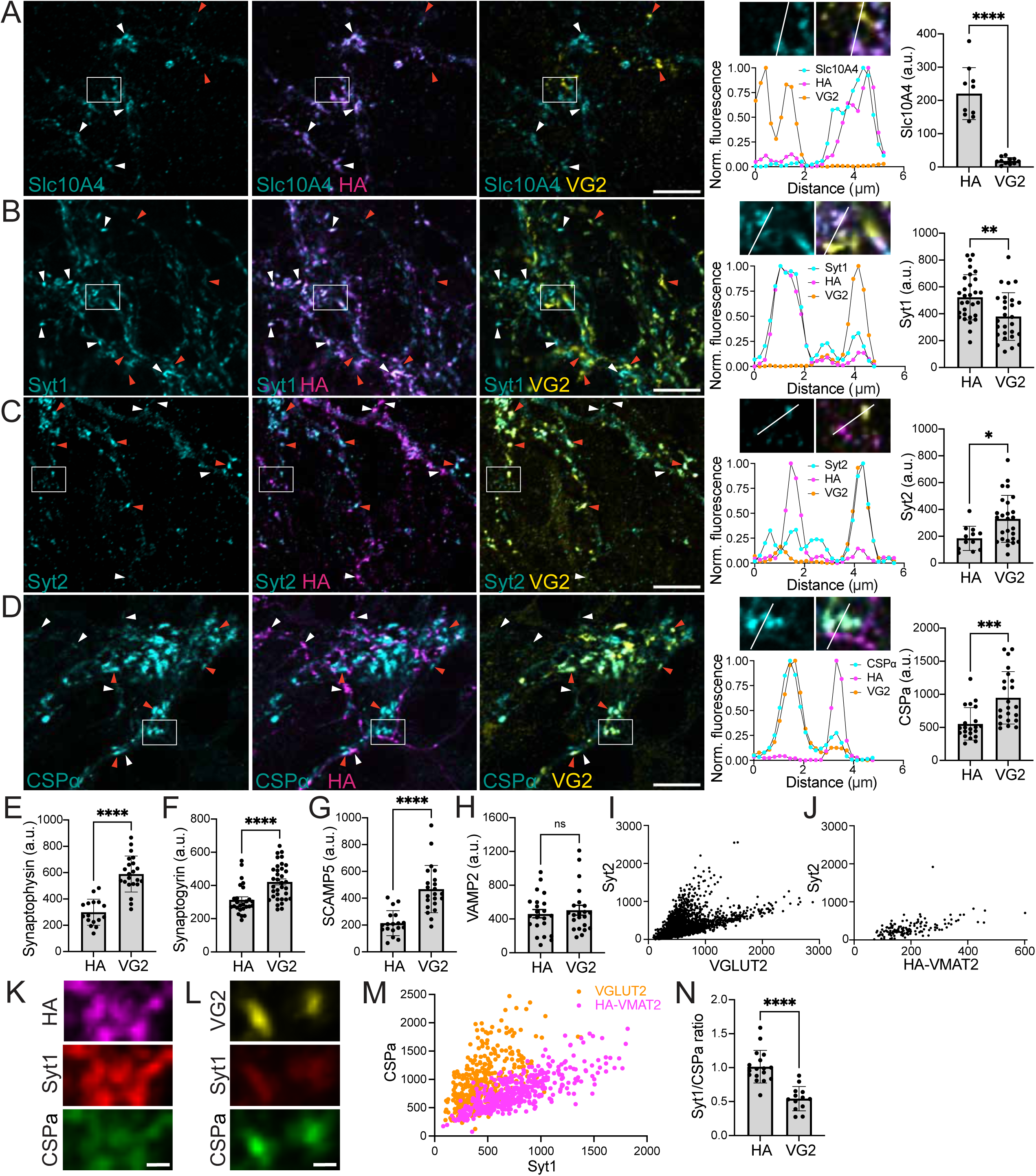
Dopamine and glutamate release sites in primary midbrain neurons differ in composition. **A-D**. Confocal images of midbrain cultures from HA-VMAT2 KI mice showing the distribution of Slc10A4 (A), syt1 (B), syt2 (C) and CSPα (D) in VMAT2 and VGLUT2 varicosities and the quantification of mean fluorescence intensity of the proteins of interest in the two types of release sites. White arrowheads point to VMAT2 and orange arrowheads to VGLUT2 varicosities. Scale bars = 10 µm. Inset shows magnification of the area indicated by the white box and a line ROI with corresponding fluorescence profiles in three channels shown below. **E-H**. Quantification of mean fluorescence intensity of synaptophysin (E), synaptogyrin1 (F), SCAMP5 (G) and VAMP2 (H) in DA release sites identified as HA-VMAT2 clusters in TH+ cells and in VGLUT2-positive glutamate release sites. Each data point represents the mean intensity of the signal for proteins of interest in all ROIs per image. **I-J**. Scatter plot of syt2 and VGLUT2 (I) and syt2 and HA-VMAT2 (J) fluorescence intensity in individual release sites. **K-N**. Presynaptic varicosities stained for HA-VMAT2, synaptotagmin1 and CSPα (K) or VGLUT2, synaptotagmin1 and CSPα (L). Scale bar = 1 µm. Scatter plot of intensities of CSPα and syt1 signal quantified in individual HA-VMAT2 or VGLUT2 varicosities (M) and quantification of syt1/CSPα ratio averaged per image (N). Data indicate mean ± SEM in all panels. * < 0.05, ** <0.01, *** < 0.001, **** <0.0001, by unpaired t-test (A, B, C, E, G, N), or Mann-Whitney test (D, F, H).

To directly test whether dopamine and glutamate release sites differ in the copy number of SV proteins, we quantified both VMAT2-enriched syt1 and VGLUT2-enriched CSPα in the same ROIs (Figure 2K-N, S5). The ratio of syt1/CSPα is significantly higher in VMAT2^+^ varicosities indicating that individual dopamine release sites contain more syt1 and less CSPα than VGLUT2 terminals (Figure 2M, 3N). These results support the differences in SV composition identified by mass spectrometry and show that possible differences in SV cluster size in dopamine versus glutamate neurons^42^ cannot alone explain differences in SV protein abundance in VGLUT2 versus VMAT2 release sites.

### The composition of synaptic vesicles differs at dopamine and glutamate release sites in the striatum

To test the differential expression of SV proteins at dopamine and glutamate release sites within the intact neural circuitry, we stained for candidate proteins in brain slices from HA-VMAT2 KI mice and quantified their abundance in VMAT2^+^ and VGLUT2^+^ varicosities (Figure 3, Figure S6). We focused on the striatum since it receives both dense dopamine projections from the midbrain and VGLUT2^+^ input from the thalamus. We acquired z-stacks and used automatic segmentation to generate three-dimensional (3D) VGLUT2 and VMAT2 surfaces followed by quantification of the mean fluorescence intensity for the proteins of interest within the same image (Figure 3A). As predicted by the proteomics results and confirmed in primary culture, Slc10A4 is enriched at dopamine release sites and colocalizes with VMAT2 (Figure 3B, 3C). If combined with TH staining to differentiate dopamine release sites from cholinergic terminals, which also express this transporter^56^, Slc10A4 can serve as a reliable marker for dopamine release sites. CSPα (Figure 3D, 3E), SCAMP5 (Figure S6A, S6B) and synaptophysin (Figure S6C, S6D) are more abundant in VGLUT2^+^ varicosities. As in primary cultures, the mean intensity of VAMP2 signal was comparable in VGLUT2^+^ and VMAT2^+^ surfaces (Figure S6E, S6F).

**Figure 3.**
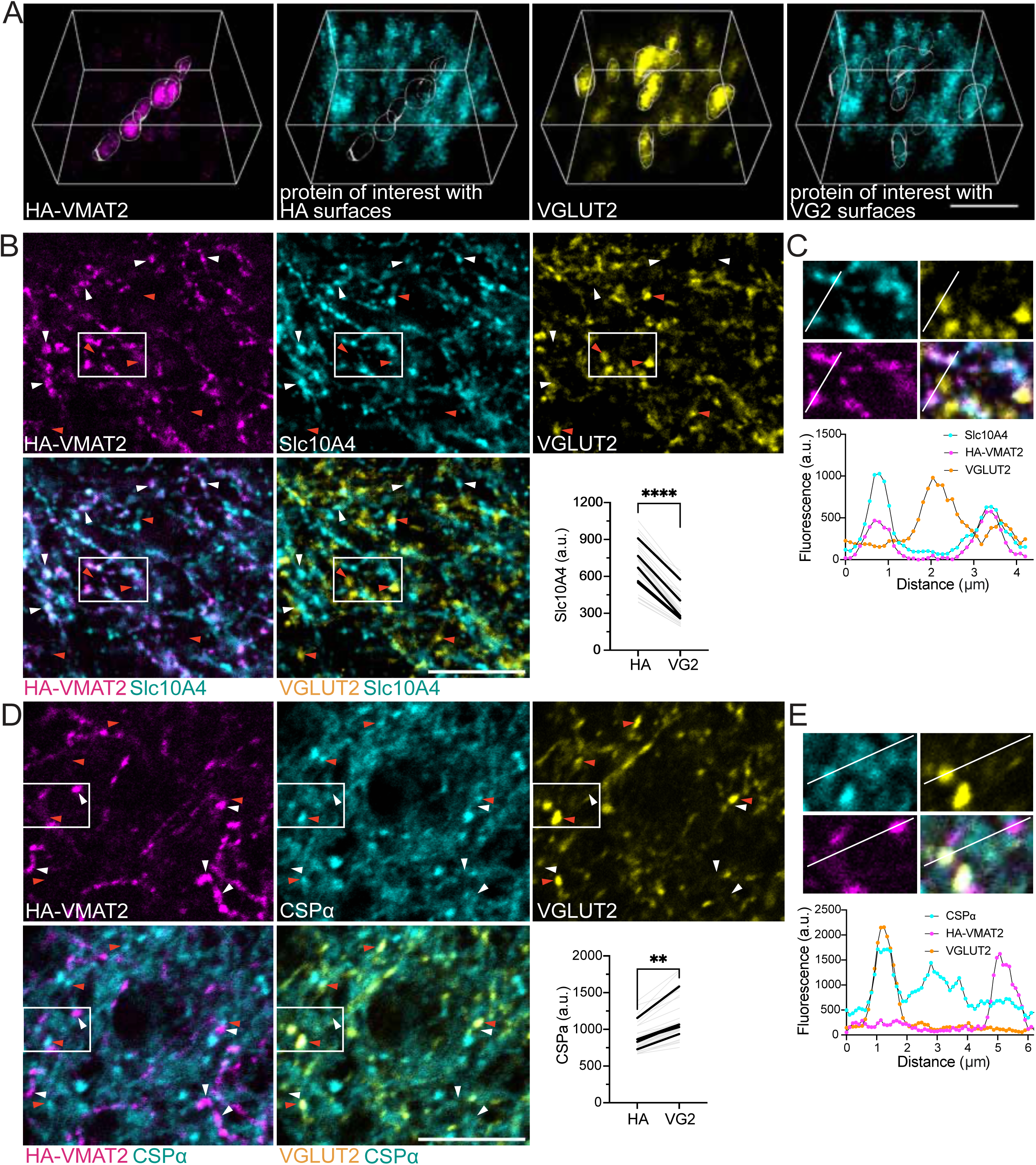
Dopamine release sites and VGLUT2^+^ glutamate terminals in the striatum differ in the abundance of SV proteins. **A.** Example of 3D surfaces generated in a striatal z-stack of a staining for HA-VMAT2 (magenta) or VGLUT2 (yellow). The same surfaces are transferred to the channel showing the staining for a protein of interest (cyan) for quantification of its the mean intensity in HA-VMAT2 and VGLUT2 surfaces. Scale bar = 4 µm. **B-C.** B. Confocal image of a single plane in a striatal slice from a HA-VMAT2 KI mouse stained for HA-VMAT2, Slc10A4 and VGLUT2 together with quantification of mean Slc10A4 intensity in 3D surfaces generated based on HA-VMAT2 or VGLUT2 signal. Gray lines show mean values of varicosities per image, black lines indicate the mean values per mouse. **** <0.0001 by paired t-test on mouse averages. Scale bar = 10 µm. White arrowheads point to HA-VMAT2 and orange arrowheads to a VGLUT2 varicosities. C. Magnified view of the area highlighted by white box in B and plot profile of fluorescence intensities in three channels along the line ROI. **D-E.** D. Confocal image of a single plane in a striatal slice from a HA-VMAT2 KI mouse stained for HA-VMAT2, CSPα and VGLUT2 together with quantification of mean CSPα intensity in 3D surfaces generated based on HA-VMAT2 or VGLUT2 signal. Gray lines show mean values of varicosities per image, black lines indicate the mean values per mouse. ** = 0.006 by paired t-test on mouse averages. Scale bar = 10 µm. White arrowheads point to HA-VMAT2 and orange arrowheads to a VGLUT2 varicosities. Magnified view of the area highlighted by white box in D and plot profile of fluorescence intensities in three channels along the line ROI.

Experiments in primary cultures indicated that syt2 is present in only a subset of VGLUT2^+^ neurons (Figure 2I). We stained for syt2 in multiple brain regions that receive VGLUT2^+^ projections to determine whether VGLUT2^+^ glutamate neurons in different structures differ in expression of syt2 (Figure 4). We focused on the striatum, CA2 region of the hippocampus, and the posterior thalamic nucleus (PO), dorsal lateral geniculate nucleus (DLG) and ventral posterior nucleus (VPN)^57^. We observed HA-VMAT2^+^ fibers in all these structures, enabling us to compare the fraction of VMAT2^+^ and VGLUT2^+^ varicosities that contain syt2 as well as to compare syt2 intensity at the two classes of release site. VMAT2^+^ and VGLUT2^+^ varicosities were considered syt2^+^ if 40% of their volume overlapped with syt2 labeling (Figure 4A). Since none of the three presynaptic proteins are expressed by all neurons, the labeling density in all brain regions was sufficiently low to assess the extent of overlap without significant contribution of signal from adjacent structures (Figure 4B, 4C). We find that although syt2 signal is very low in VGLUT2^+^ terminals of the striatum and in region CA2 of the hippocampus, it is abundant at the majority of VGLUT2^+^ release sites in thalamic nuclei and absent from the majority of VMAT2^+^ varicosities in all structures (Figure 4B-D). When we quantified the mean intensity of syt2 signal in monoamine and glutamate release sites across all regions, we find that this calcium sensor is significantly more abundant in VGLUT2 varicosities in the DLG that contains the large VGLUT2^+^ terminals of retinal ganglion cells^58^ (Figure 4B, 4C, 4E) and in the VPN where VGLUT2-dependent glutamate release relays sensory information from the spinal cord and brain stem^59^ (Figure 4E). Mean syt2 intensity was not significantly different between VMAT2^+^ and VGLUT2^+^ varicosities in the striatum, CA2 and PO. These results confirm the differences in composition of VMAT2^+^ and VGLUT2^+^ SVs identified by the proteomics.

**Figure 4.**
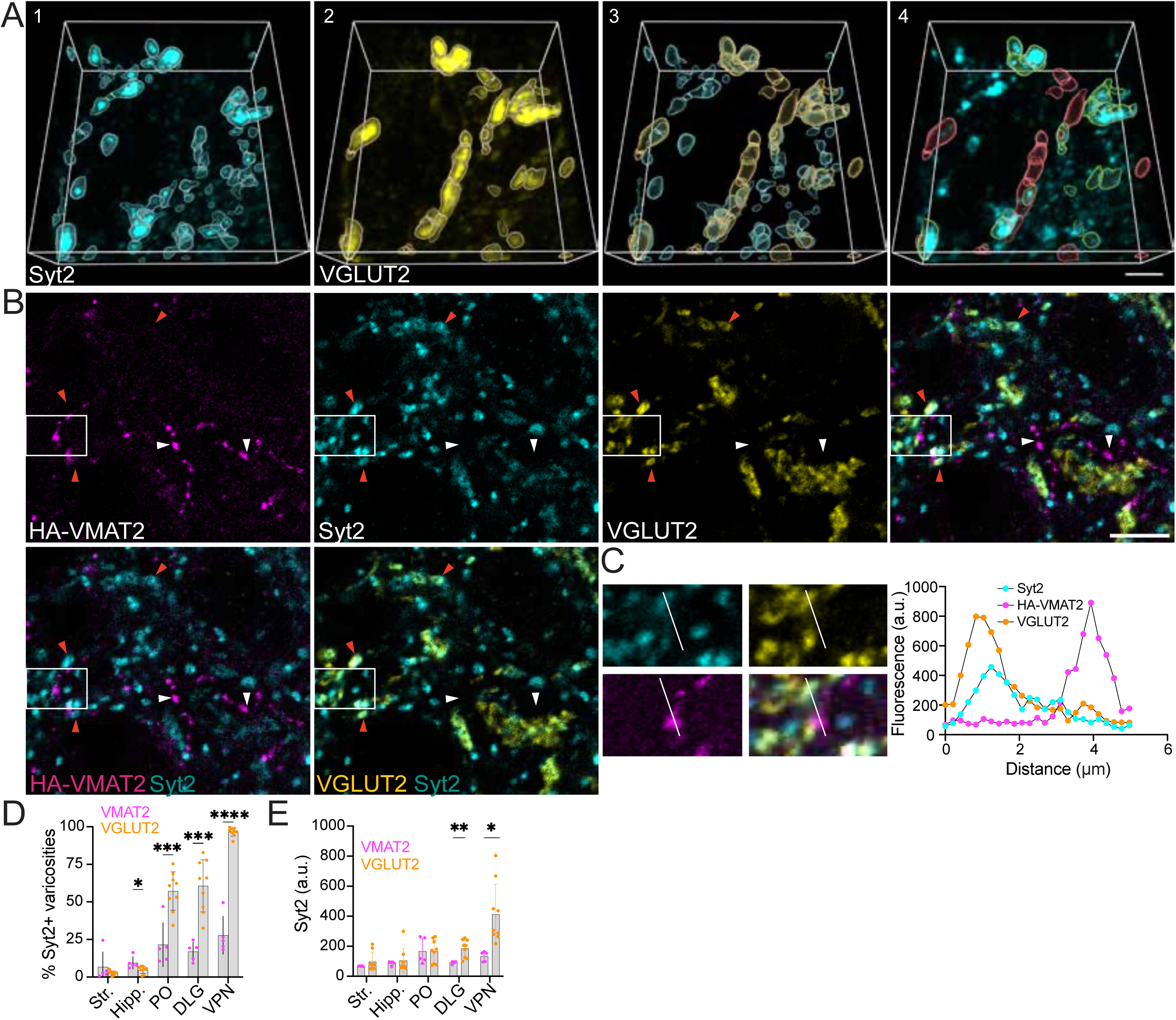
Expression of syt2 reveals heterogeneity among VGLUT2+ glutamate neurons. A. Example of 3D surfaces generated in a z-stack based on staining for syt2 (cyan, panel 1) or VGLUT2 (yellow, panel 2). Panel 3 shows syt2 and VGLUT2 surfaces in the same 3D section. In panel 4, syt2-positive VGLUT2 surfaces are shown in green and syt2-negative VGLUT2 surfaces are shown in red. A varicosity is considered syt2-positive if >40% of its volume overlaps with syt2 signal. Scale bar = 4 µm. B. A single plane of a confocal image of a fixed brain slice from a HA-VMAT2 KI mouse acquired in the lateral geniculate nucleus (DLG). The slice is stained for HA-VMAT2, syt2 and VGLUT2. White arrowheads point to VMAT2 and orange arrowheads to a VGLUT2 varicosities. Scale bar = 3 µm. C. Magnified view of the area highlighted by white box in B and plot profile of fluorescence intensities in three channels along the line ROI. D. Percentage of VGLUT2 and HA-VMAT2 varicosities that contain syt2 signal in different brain regions. Each data point represents the percentage of syt2-positive varicosities per z-stack. Data indicate mean ± SEM. * = 0.0175, *** < 0.001, **** < 0.0001 by two-tailed unpaired t-test. E. Quantification of mean syt2 intensity in HA-VMAT2 and VGLUT2 surfaces across brain regions. Each data point represents an analyzed z-stack. Data indicate mean ± SEM. * = 0.0105, ** = 0.0047 by two-tailed unpaired t-test. Str. – striatum, Hipp. – CA2 region of the hippocampus, PO – posterior complex, VPN – ventral posteromedial nucleus.

### SV protein composition of VGLUT2+ release sites in dopamine neurons differs from VMAT2-only varicosities

The corelease of glutamate by midbrain dopamine neurons raises the possibility that SVs that store dopamine and glutamate may differ within the same population of neurons. We previously found that the exocytosis and recycling of VGLUT2^+^ SVs in cultured dopamine neurons occurs with faster kinetics than VMAT2^+^ SVs^22^, indicating differences in the mode of release in the same cell. Glutamate release from coreleasing neurons also differs from dopamine release in short-term plasticity^22^. If VGLUT2^+^ and VMAT2^+^ SVs in dopamine neurons indeed differ in protein content, it should result in altered presynaptic composition in VGLUT2^+^ release sites in dopamine neurons compared to VMAT2-only sites.

The corelease of glutamate with dopamine reflects the expression of VGLUT2 by a subset of midbrain dopamine neurons^32, 33^. To investigate the localization of VMAT2 and VGLUT2 in these cells, we again took advantage of midbrain cultures from HA-VMAT2 mice (Figure 5). We find that within TH^+^ neurons that label for both VMAT2 and VGUT2, most release sites contain detectable signal for both transporters, even though the relative intensities vary (Figure 5A-C). While VMAT2 signal is comparable in release sites of VMAT2-only and VMAT2^+^/VGLUT2^+^ neurons, VGLUT2 abundance in presynaptic varicosities of cultured VMAT2^+^/VGLUT2^+^ neurons is significantly lower than in glutamate-only cells (Figure 5D). This suggests a lower VGLUT2 copy number per vesicle or its presence on a subset of vesicles within coreleasing terminals.

**Figure 5.**
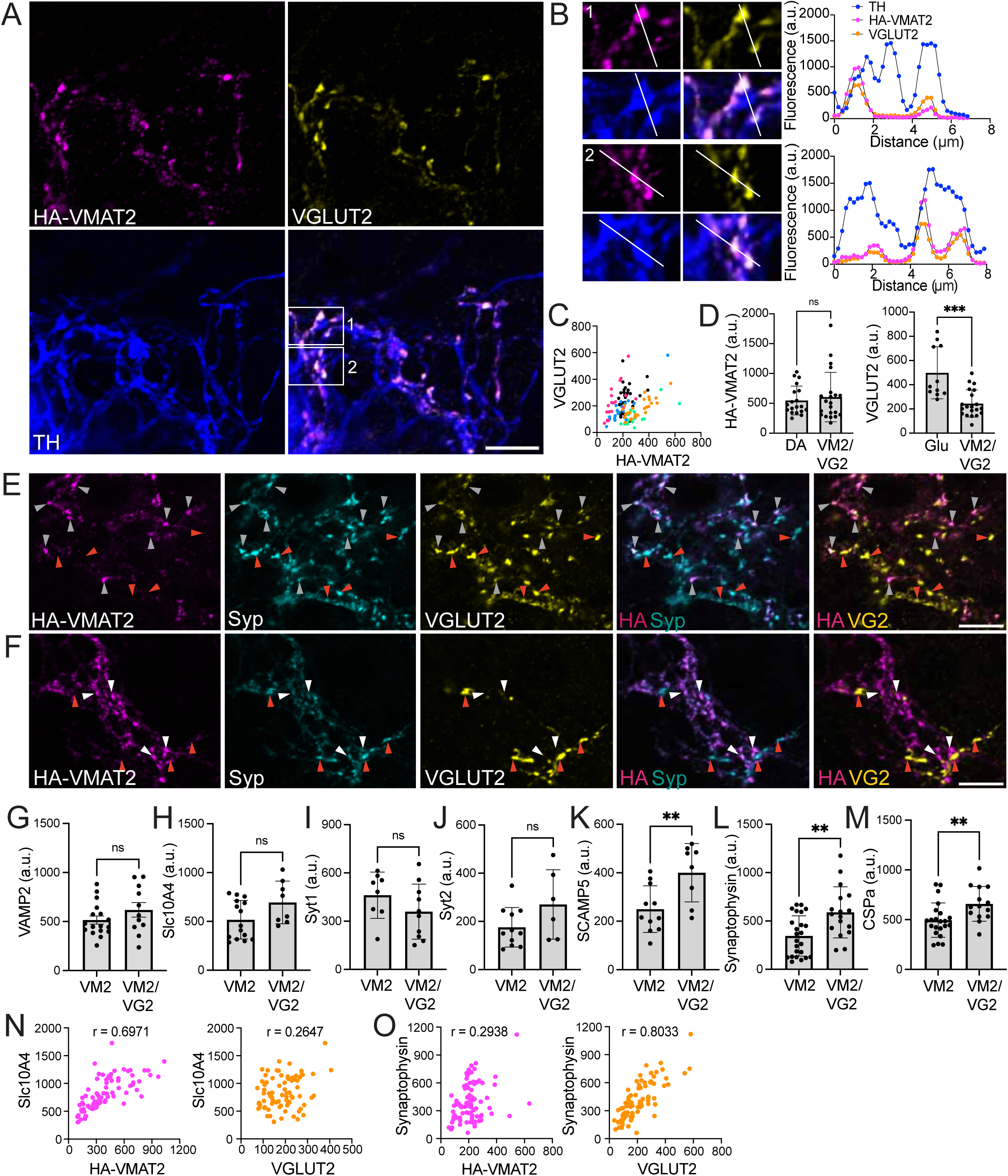
SV protein composition of VMAT2-only versus VMAT2^+^/VGLUT2^+^ release sites. **A.** Confocal image of a VGLUT2-positive DA neuron in a primary midbrain culture from HA-VMAT2 KI mice. Scale bar = 10 µm. **B.** Magnified view of areas indicated by white boxes in A and plot profile of HA-VMAT2, VGLUT2 and TH signal along the line ROI showing overlapping VGLUT2 and HA-VMAT2 peaks in a DA neuron. **C.** Scatter plot of HA-VMAT2 and VGLUT2 signal in individual presynaptic varicosities from one data set. Each color represents varicosities from one image (likely corresponding to a single cell). **D.** Quantification of HA-VMAT2 signal in the presynaptic terminals of dopamine-only versus VMAT2^+^/VGLUT2^+^ neurons and VGLUT2 signal in glutamate-only versus VMAT2^+^/VGLUT2^+^ neurons. Each data point represents mean of individual ROIs per image. Data indicate mean ± SEM. *** = 0.0006 by Mann-Whitney test. **E.** Confocal image of a coreleasing neuron and adjacent glutamate-only terminals stained for TH (not shown), synaptophysin, HA-VMAT2 and VGLUT2. Gray arrowheads point to VGLUT2^+^/VMAT2^+^ varicosities and orange arrowheads to VGLUT2-only terminals. Note that all HA-VMAT2 varicosities have detectable VGLUT2 signal. Scale bar = 10 µm. **F.** Confocal image of a dopamine-only neuron and an adjacent glutamate-only neuron stained for TH (not shown), synaptophysin, HA-VMAT2 and VGLUT2. White arrowheads point to VMAT2 terminals and orange arrowheads to VGLUT2 terminals. Scale bar = 10 µm. **G-M**. Quantification of VAMP2 (G), Slc10A4 (H), syt1 (I), syt2 (J), SCAMP5 (K), synaptophysin (L) and CSPα (M) in presynaptic terminals of dopamine-only or VMAT2^+^/VGLUT2^+^ neurons. Data indicate mean ± SEM. Each data point represents mean intensity of the signal for a protein of interest in all presynaptic varicosities per image. Data indicate mean ± SEM. ** <0.01 by unpaired t-test (K, M) or Mann-Whitney test (L). **N**. Scatterplots showing correlation between Slc10A4 and HA-VMAT2 (magenta) or Slc10A4 and VGLUT2 (orange) signal in the same set of presynaptic terminals that contain both VGLUT2 and VMAT2. **O**. Scatterplots showing correlation between synaptophysin and HA-VMAT2 (magenta) or synaptophysin and VGLUT2 (orange) signal in the same set of presynaptic terminals that contain both VGLUT2 and VMAT2.

We also determined whether VMAT2^+^/VGLUT2^+^ sites differ from VMAT2-only varicosities in the expression of other SV proteins. We find that VMAT2-enriched Slc10A4 and equally distributed VAMP2 are present at similar levels in VMAT2-only and VMAT2^+^/VGLUT2^+^ boutons (Figure 5G-H). Likewise, the abundance of SV calcium sensors syt1 and syt2 is not affected by the presence of VGLUT2 at VMAT2 sites (Figure 5I-J). However, CSPα, SCAMP5 and synaptophysin are significantly more abundant at VMAT2^+^/VGLUT2^+^ boutons compared to VMAT2-only SV clusters (Figure 5E, 5F, 5K-M). Furthermore, we find that the abundance of Slc10A4 measured in individual release sites of coreleasing neurons correlates better with VMAT2 than VGLUT2 abundance at the same varicosities (Figure 5N). Conversely, synaptophysin signal correlates with VGLUT2 expression (Figure 5O) supporting the preferential targeting of synaptophysin to VGLUT2 SVs. Altogether, we find that VGLUT2^+^ presynaptic varicosities in dopamine neurons resemble typical glutamate release sites pointing to a distinct composition of VGLUT2 SVs within dopamine neurons.

### SVs containing VGLUT2 and VMAT2 segregate within presynaptic varicosities of dopamine/glutamate coreleasing neurons

VMAT2 and VGLUT2 colocalize at the majority of release sites made by cultured dopamine neurons, but previous work using heterologous reporters has suggested that they may localize to distinct SVs at these sites^22^. We have now addressed this question using endogenous proteins. Since classical confocal microscopy does not allow us to resolve SVs within a single presynaptic bouton, we thus took advantage of the Nikon Spatial Array Confocal (NSPARC) technology to access enhanced spatial resolution compared to classical confocal detection systems^60^. We focused on individual presynaptic varicosities that contain TH, HA-VMAT2 and VGLUT2 to evaluate the degree of colocalization between VMAT2 and VGLUT2 signal within the release site and to investigate the colocalization of the signal for the transporters and differentially expressed SV proteins (Figure 6). First, we compared the colocalization of HA-VMAT2 and VGLUT2 (Figure 6B, 6C), using HA signal stained with two secondary antibodies coupled to different fluorophores as control (Figure 6A). As expected, HA signal detected in two channels shows a high degree of colocalization. We also detect substantial overlap between HA-VMAT2 and VGLUT2 signal, however, it is significantly lower compared to the control condition comparing HA signal in two channels (Figure 6A). Since we detect a partial segregation of VMAT2 and VGLUT2 into different structures within presynaptic varicosities, we compared the correlation of VGLUT2-enriched synaptophysin and VMAT2-enriched syt1 with the signal for the two transporters. We find that synaptophysin fluorescence correlates more with VGLUT2 than VMAT2 in presynaptic terminals that contain both transporters (Figure 6B) while syt1 signal displays similar correlation with both VGLUT2 and VMAT2 (Figure 6C). These results suggest that VGLUT2 targets to an SV population with distinct composition.

**Figure 6.**
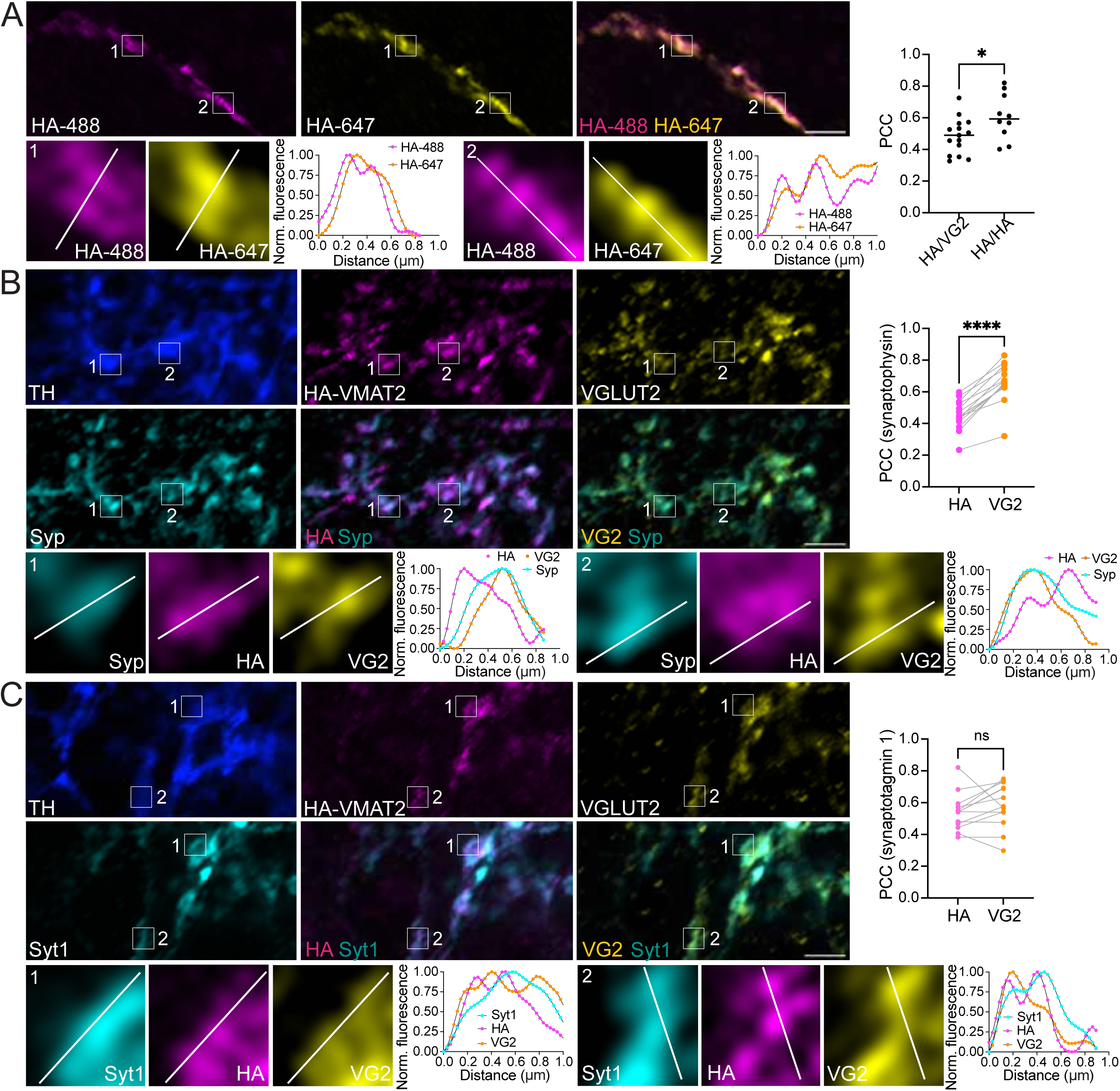
Synaptophysin is co-enriched with VGLUT2 in dopamine neuron presynaptic varicosities that contain both VMAT2 and VGLUT2. **A.** Dopamine neuron from an HA-VMAT2 culture double stained for HA-VMAT2 using secondary antibodies coupled with Alexa 488 and Alexa 647 fluorophores. Scale bar = 2 µm. Quantification shows Pearson’s correlation coefficient (PCC) in individual varicosities for HA imaged in two channels versus for HA and VGLUT2 measured in data sets shown in B and C. Data indicate mean ± SEM. Each data point represents the mean of individual varicosities per image. * = 0.0265 by unpaired t-test. N = 10 HA/HA and 15 HA/VGLUT2 coverslips from two independent cultures. Insets show magnification of areas indicated with white boxes with line ROIs and fluorescence profiles in HA-488 and HA-647 channels. **B.** VGLUT2^+^ DA neuron from an HA-VMAT2 midbrain culture stained for TH, HA, VGLUT2 and synaptophysin (syp). Scale bar = 2 µm. Quantification shows Pearson’s correlation coefficient (PCC) for HA and synaptophysin versus VGLUT2 and synaptophysin quantified in varicosities that contain all three proteins. Data indicate mean ± SEM. Each data point represents the mean of individual varicosities per image. **** < 0.0001 by paired t-test. N = 14 images from two independent cultures. Insets show magnification of areas indicated with white boxes with line ROIs and fluorescence profiles in HA, VGLUT2 and synaptophysin channels. **C.** VGLUT2^+^ DA neuron from an HA-VMAT2 midbrain culture stained for TH, HA, VGLUT2 and synaptotagmin1 (syt1). Scale bar = 2 µm. Quantification shows Pearson’s correlation coefficient (PCC) for HA and synaptotagmin1 versus VGLUT2 and synaptotagmin1 quantified in varicosities that contain all three proteins. Data indicate mean ± SEM. Each data point represents the mean of individual varicosities per image. P = 0.2181 by paired t-test. N = 13 images from two independent cultures. Insets show magnification of areas indicated with white boxes with line ROIs and fluorescence profiles in HA, VGLUT2 and synaptotagmin1 channels.

### Loss of SCAMP5 selectively impairs the synaptic recycling of VGLUT2 vesicles

If VGLUT2 and VMAT2 SVs differ in composition even in the same neuronal population, then elimination of a differentially expressed protein should preferentially affect the synaptic recycling of vesicles expressing more of that protein. We focused on SCAMP5 as we have validated its significantly higher abundance on VGLUT2 vesicles and previous studies have shown that its loss of function affects the SV cycle and neurotransmitter release^61, 62^. To address its role in the release of VMAT2^+^ versus VGLUT2^+^ vesicles, we took advantage of the pHluorin reporter that enables monitoring the properties of the SV cycle when fused to luminal domains of SV proteins. PHluorin fluorescence is quenched in resting, acidic SVs but increases as SV fusion exposes the vesicle lumen to the neutral extracellular environment and decreases with endocytosis and vesicle reacidification^63^. We have previously shown that pHluorin-tagged VMAT2 reliably targets to AP-3-dependent SVs in both dopamine and hippocampal neurons whereas VGLUT2-pHluorin is excluded from these vesicles^22, 27^. To test whether the elimination of SCAMP5 has differential effects on these SV pools, we used shRNA to knock down (KD) SCAMP5 expression (Figure 7). Lentiviral delivery of a KD construct targeting SCAMP5 results in an efficient elimination of the SCAMP5 protein from hippocampal neurons without affecting neuronal morphology or the abundance of the SV calcium sensor syt1 (Figure 7A). To test the role of SCAMP5 in the VGLUT2^+^ SV population, we infected neuronal cultures with VGLUT2-pHluorin together with a control scrambled construct or SCAMP5 KD alone or in combination with an shRNA resistant rescue construct. Neurons were stimulated for 60 seconds at 10 Hz. We find that the recycling of VGLUT2 SVs is reduced in the KD condition compared to a scrambled control but is restored to control levels if SCAMP5 expression is rescued in the KD background (Figure 7B). The synaptic recycling of AP-3-dependent SVs targeted by VMAT2-pHluorin is however not affected by the loss of SCAMP5 (Figure 7C). These findings indicate that SCAMP5 is specifically involved in recycling of vesicles that are enriched in VGLUT2 and despite its high abundance in the cell type, does not have a detectable effect on the trafficking of the SV population targeted by VMAT2. Since the loss of AP-3 does not completely abolish VMAT2 recycling^27^, it remains possible that AP-3-independent VMAT2^+^ vesicles depend on SCAMP5 function but this effect is not detected due to the abundance of VMAT2 on AP-3 SVs.

**Figure 7.**
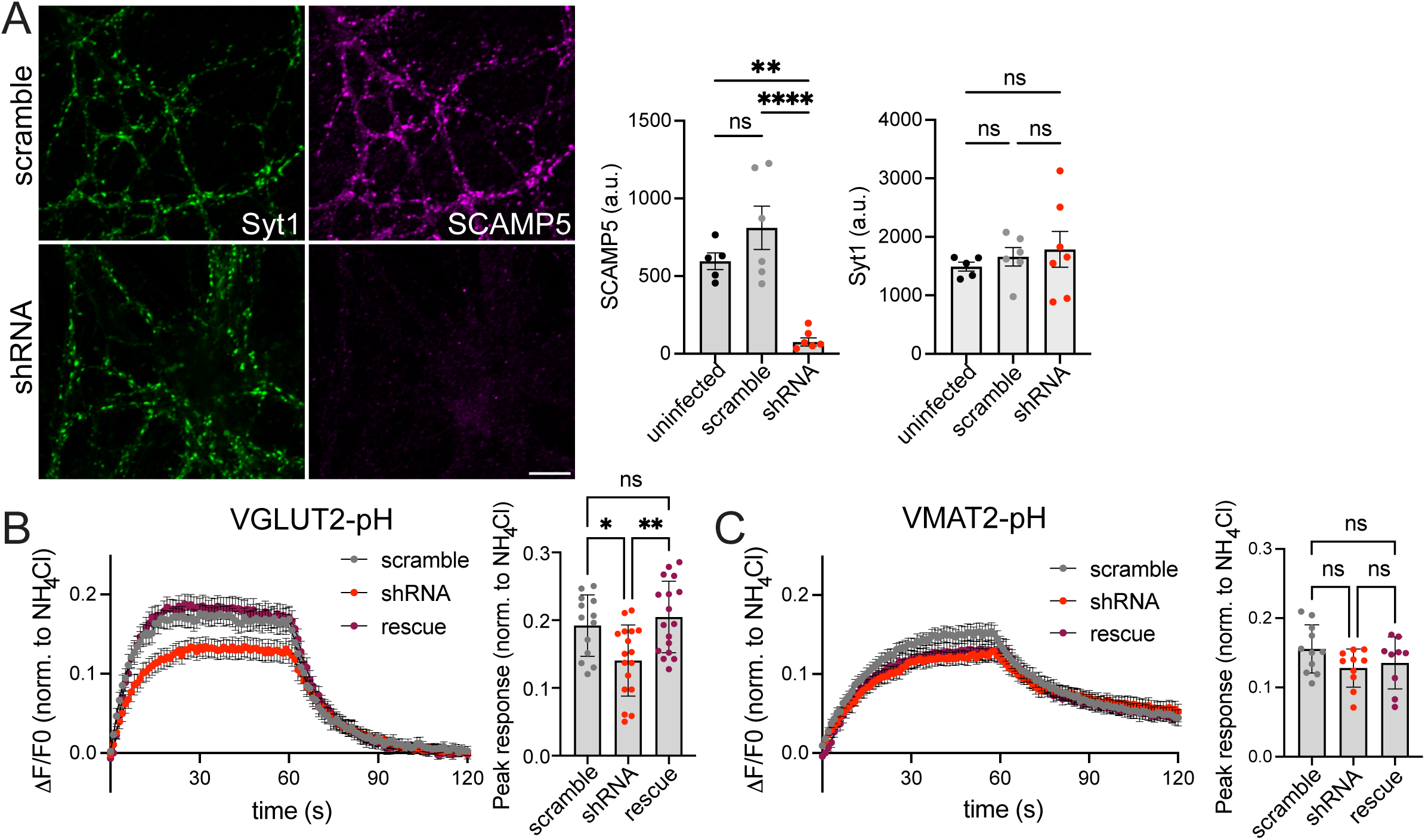
Loss of SCAMP5 impairs synaptic recycling of VGLUT2 SVs without affecting VMAT2. A. Confocal images and quantification of the fluorescence intensity of synaptotagmin1 and SCAMP5 in uninfected hippocampal neurons or cultures infected with shRNA targeting SCAMP5 or a scrambled control. Each data point represents mean presynaptic SCAMP5 or syt1 intensity per image. Data indicate mean ± SEM, ** p = 0.0021, **** p < 0.0001 by one-way ANOVA with Tukey’s multiple comparisons. n = 5 uninfected, 6 scramble and 7 shRNA coverslips. B-C. Average response curves and quantification of peak fluorescence intensity of VGLUT2-pHluorin (B) or VMAT2-pHluorin (C) to a 60s stimulation at 10 Hz. Data indicate mean ± SEM, * = 0.0225, ** = 0.017 by one-way ANOVA with Tukey’s multiple comparisons. N = 13 scramble, 17 shRNA and 17 rescue coverslips for VGLUT2-pHluorin and 11 scramble, 10 shRNA and 9 rescue coverslips for VMAT2-pHluorin from three independent cultures.

## DISCUSSION

The results show that VMAT2^+^ and VGLUT2^+^ synaptic vesicles differ in protein composition. VMAT2^+^ vesicles contain fewer canonical SV proteins than VGLUT2^+^ SVs but more of others. These findings provide a basis to understand the different properties of monoamine release and glutamate release that depends on VGLUT2.

We identify some of the same unique components of dopamine SVs that were detected in previous studies that relied on synaptosome sorting or proximity labeling^16, 26, 64, 65^. These include putative transporters Slc10A4, VAT1 and SV2c^16, 64, 65^. While their substrate in dopamine neurons and mechanism of action is not completely understood, the loss of Slc10A4 and SV2c leads to reduced dopamine uptake into SVs and impaired release^43, 66, 67^. VAT1 was recently shown to bind NADPH to function as a quinone oxidoreductase^68^. While its synaptic role in not known, its enzymatic function could contribute to alleviation of oxidative stress.

The enrichment of Slc10A4 and SV2c in VMAT2+ SVs is likely explained by their high expression in monoamine neurons and absence from many other neuronal populations^13–15^. However, we also find differences in SV proteins that are expressed in most neurons. For example, VMAT2^+^ SVs contain relatively little synaptophysin and closely related synaptogyrin1 and synaptoporin. While the primary function of this protein family is not clear, their loss of function affects both short- and long-term synaptic plasticity^69, 70^ as well as trafficking of other SV components^71–73^. Recent evidence also points to a role of this protein family in the maintenance of SV size, with larger SVs observed in the absence of synaptophysin^74, 75^. Furthermore, synaptophysin enables the expansion of filling SVs, which accelerates their fusion^75^. Low synaptophysin abundance on VMAT2^+^ vesicles could therefore affect their function as well as morphology, which is consistent with recent reports of larger and less uniform SVs in dopamine release sites^42^.

Among other canonical SV proteins, we find that VMAT2^+^ and VGLUT2^+^ vesicles differ in synaptotagmin isoforms. Dopamine release sites are enriched in the fast Ca^++^ sensor syt1 and lack the closely related fast isoform syt2 that is abundant in a subset of VGLUT2^+^ glutamate neurons that relay sensory information in a highly reliable manner. Additionally, VMAT2^+^ SVs are enriched in syt5 and VGLUT2^+^ SVs in syt12. Unlike previous studies^64, 65^, we do not find a significant enrichment of syt17 in VMAT2^+^ vesicles. However, functional data supports syt17 localization on endosomal organelles that differ from SVs^76, 77^ which could explain its enrichment in dopamine synaptosomes and axons but not SVs. Abundance of syt1 on VMAT2^+^ SVs is consistent with recent work which shows that this Ca^++^ sensor is required for most dopamine release^78, 79^. While syt1 and syt2 share high structural and functional similarity, several studies have reported faster kinetics and higher temporal precision of release mediated by syt2^47, 80^ even though this was not seen in all synapses^81, 82^. The function of syt1 and syt12 but not syt2 is also modulated by protein kinases A and C – known regulators of presynaptic plasticity^45, 46, 83^. Further investigation is therefore needed to elucidate whether the expression of different synaptotagmins contributes to the distinct kinetics and plasticity of dopamine and glutamate release.

VMAT2^+^ and VGLUT2+ SVs also differ in SCAMP isoforms. SCAMP4 and −5 are more abundant on VGLUT2^+^ vesicles and SCAMP2 is on VMAT2^+^ SVs. Consistently with its role in monoamine release sites, loss of SCAMP2 has previously been shown to impair the trafficking of the dopamine transporter DAT and serotonin transporter SERT^53, 84^. In the context of neurotransmitter release, SCAMP5 has been studied in most detail with multiple roles identified at the presynaptic terminal. Its loss of function was shown to reduce the quantal content of glutamate release by impairing presynaptic targeting of the cation/H^+^ exchanger NHE6^85^ and to enhance synaptic depression via impaired release site clearance and endocytosis^61, 62^ but it is unclear whether these roles are common to all isoforms. Direct comparison of SCAMPs in dense core vesicle exocytosis suggests that while knock-downs of all studied isoforms reduce the number of exocytotic events, they have distinct effects on the latency of exocytosis and only SCAMP1 and −5 have additional effects on transmitter uptake^86^. Differential enrichment of SCAMP isoforms on VMAT2^+^ and VGLUT2^+^ SVs is therefore likely to have distinct effects on release. Indeed, we now demonstrate that the loss of SCAMP5 impairs the release of VGLUT2^+^ SVs without affecting the vesicles targeted by VMAT2 even in the same neuronal population. These findings demonstrate that differences in SV composition confer distinct regulation to VMAT2^+^ and VGLUT2^+^ SVs, which in turn mediate the release of dopamine and glutamate.

Beyond SV membrane proteins, we also find that VMAT2^+^ vesicles contain less of the lipid-anchored neuroprotective chaperone CSPα. CSPα contributes to the correct folding of the release machinery and its loss of function leads to a rapid presynaptic degeneration^87^, while a reduced copy number impairs synaptic function more slowly^88^. The low expression of CSPα at dopamine release sites could therefore increase dopamine neuron susceptibility to protein misfolding which might in turn contribute to their vulnerability in Parkinson’s disease. In addition to SV proteins, we find that dynactin and dynein subunits are preferentially associated with VMAT2^+^ SVs, as reported before^16^. Association with the dynein motor could reflect anterograde trafficking of VMAT2 in dendrites or a more dynamic nature of diffusely distributed SVs in dopamine axons^18, 41^, which in turn could affect their availability for release^89, 90^.

Differences in the abundance of SV proteins that are widely expressed across neurons suggests that the formation of VMAT2+ and VGLUT2+ SVs depends on distinct mechanisms that differ in efficiency or selectivity of protein recruitment. Indeed, the reliance of VMAT2 but not VGLUT2 on the AP-3 pathway is likely to contribute to differences in SV composition and release^12, 22, 27^. Previous analysis of whole brain SVs in WT versus AP-3 knock-out mice confirmed the reduced expression of known AP-3 client proteins, including VAMP7 and identified new SV proteins that depend on AP-3 function, including phospholipid flippase ATP8a1 and Slc35g2 – a transporter with unknown function^30, 91, 92^. While the abundance of most classical SV proteins was not found to be changed^30^, AP-3-dependent vesicles only constitute a small fraction of the total SV pool at most synapses, making it challenging to detect changes in the abundance of proteins that do not rely exclusively on AP-3 function for SV targeting^31, 93^. We expected to find enrichment of known AP3-dependent proteins in VMAT2+ SVs, however, while Slc35g2 appears abundant in these vesicles, VAMP7 and ATP8A1 do not differ in enrichment in the two SV populations and surprisingly, AP-3-dependent zinc transporter ZnT3 is enriched in the VGLUT2^+^ population. ZnT3 targeting is partially explained by its lack of expression in dopamine neurons and presence in a small subset of VGLUT2^+^ glutamate neurons^14^, however, it remains unclear how it is packaged onto vesicles together with VGLUT2 that does not rely on AP-3. Additional work is also needed to elucidate the localization of AP-3-dependent proteins VAMP7 and VMAT2 in the presynaptic terminal. Indeed, VAMP7 has established roles in endolysosomal organelles^94–96^ in addition to SVs^30, 31, 91, 93^ and could therefore reside in a partially separate vesicle population at the release site^97^. Among proteins enriched in VGLUT2^+^ SVs, SCAMP5 is known to interact with AP-2 and this interaction is required for its role in release sites clearance^62^. Synaptophysin has been shown not to depend on AP-3^92, 94^. Furthermore, its association with other SV proteins can affect their sorting^72, 73^ and can therefore contribute to generating differences in the composition of VMAT2^+^ and VGLUT2^+^ SVs. Indeed, the distribution of known AP-3 cargos in the two vesicle populations shows that reliance on distinct cargo adaptors is likely not sufficient to generate the observed differences in protein composition. Indeed, it is likely that cell type-specific differences in gene expression, differential reliance on adaptors as well as sorting by association with other SV proteins all contribute to establishing the distinct composition of VMAT2^+^ and VGLUT2^+^ SVs^98^.

A significant role of distinct trafficking mechanisms is further highlighted by the analysis of VGLUT2^+^ dopamine neurons. We find that while HA-VMAT2 and VGLUT2 colocalize at most release sites in cultured dopamine/glutamate cells, they often differ in abundance. VGLUT2^+^ release sites in dopamine neurons also differ in SV protein composition from VMAT2-only sites suggesting a distinct composition of VGLUT2^+^ SVs. While the analysis of VMAT2-only and VMAT2^+^/VGLUT^+^ sites mostly compares different cells and could reflect differences in gene expression between coreleasing and dopamine-only neurons, super-resolution imaging reveals a partial segregation of VMAT2 and VGLUT2 as well as co-enriched SV proteins in individual presynaptic varicosities. These findings show that VMAT2 and VGLUT2 are packaged into biochemically distinct vesicles in the same cell and even presynaptic terminal. The specific effect of SCAMP5 KD on VGLUT2^+^ vesicles further demonstrates that differences in SV composition translate into distinct regulation of release.

Targeting of VMAT2 and VGLUT2 to distinct vesicles is compatible with previous work, which demonstrated the different kinetics and frequency dependence of dopamine and glutamate release from coreleasing neurons^22^. However, glutamate corelease was also shown to affect dopamine packaging into SVs via vesicular synergy, which would require at least a partial coexistence of VMAT2 and VGLUT2 on the same vesicle^32^. Indeed, recent findings show that while AP-3 is important for generating VMAT2^+^ vesicles, the loss of this adaptor does not completely abolish dopamine release^27^. This suggests that some VMAT2 localizes to AP-3-independent SVs, which likely also contain VGLUT2. It is therefore possible that colocalization of the transporters on the AP-3-independent SV pool drives vesicular synergy observed between the two transmitters^32^.

In summary, we elucidate the distinct molecular composition of monoamine and VGLUT2^+^ glutamate SVs. The differential role of SCAMP5 illustrates the functional significance of these differences. Other differentially expressed SV proteins presumably contribute to differences in dopamine and glutamate release.

**Supplementary figure 1.**
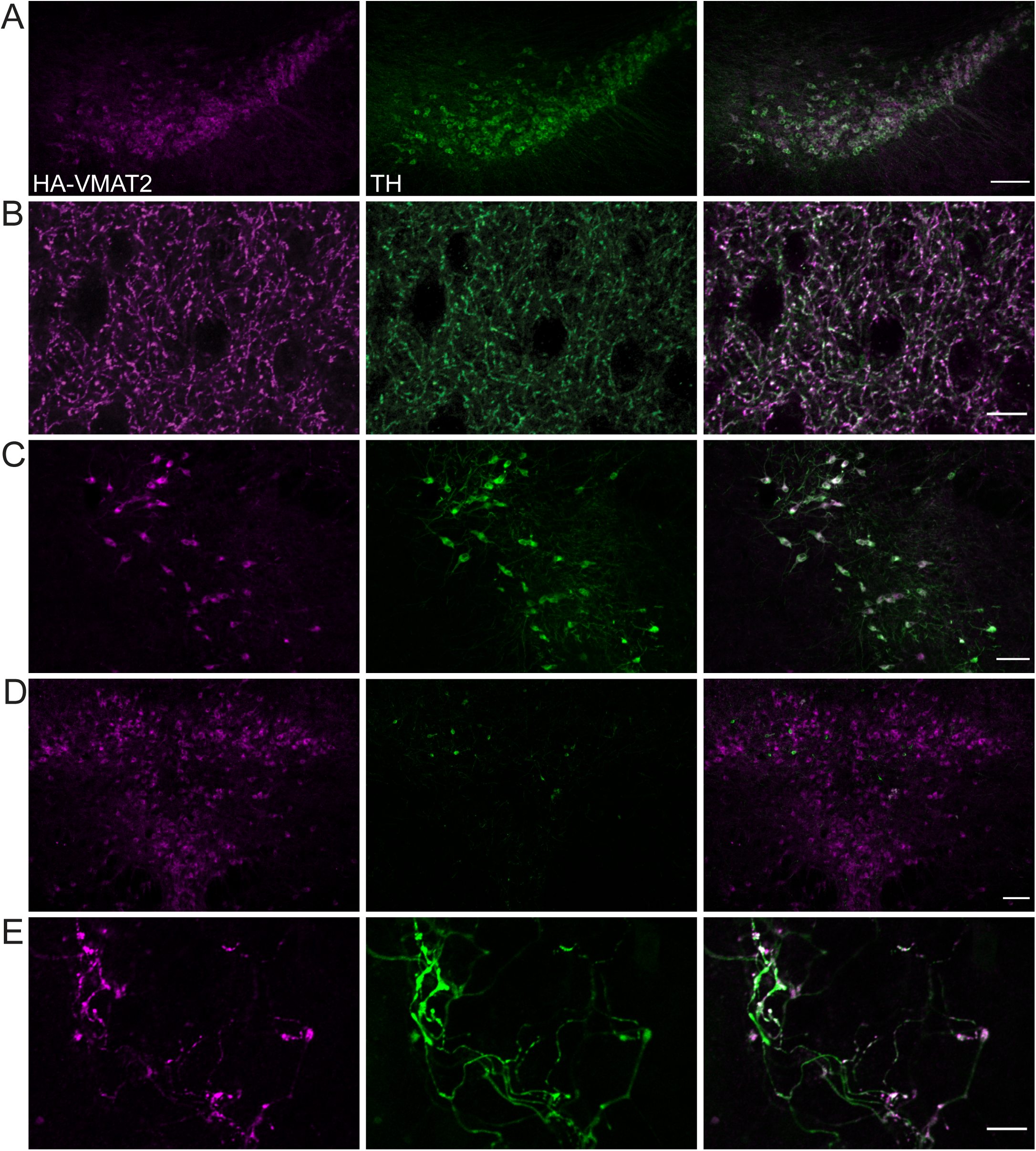
HA-VMAT2 is localized in cell bodies and neuronal processes of monoamine neurons. A-D. Representative confocal images of HA-VMAT2 KI brain slices from the midbrain (A), striatum (B), locus coeruleus (C) and the raphe nucleus (D) stained for HA (magenta) and tyrosine hydroxylase (TH, green). Scale bar = 10 µm in all images. E. Representative confocal image of a midbrain culture form HA-VMAT2 mice. Cultured neuron is stained for HA and TH. Scale bar = 10 µm.

**Supplementary figure 2.**
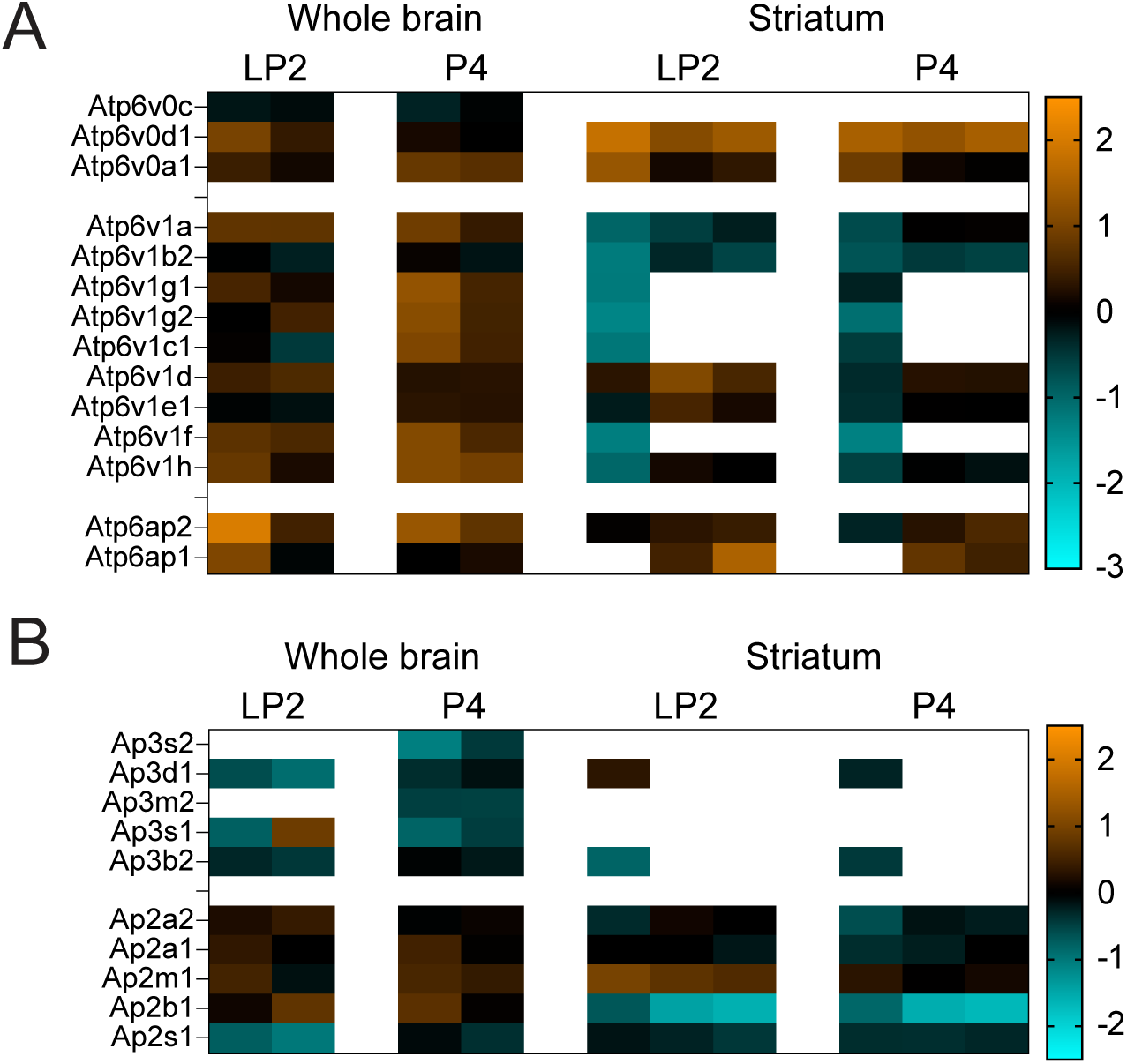
Abundance of V-ATPase, AP-2 and AP-3 subunits in VMAT2+ and VGLUT2+ SVs. A. Subunits of the v-ATPase quantified on VMAT2 and VGLUT2 SVs isolated from whole brain or striatal samples from the HA-VMAT2 KI mouse. Data was normalized to the amount of total protein in each sample and is presented as a heat map of log2 values of VGLUT2/VMAT2 ratio. Each column represents an independent experiment. B. Subunits of AP-2 and AP-3 quantified on VMAT2 and VGLUT2 SVs isolated from whole brain or striatal samples from the HA-VMAT2 KI mouse. Data was normalized to the amount of total protein in each sample and is presented as a heat map of log2 values of VGLUT2/VMAT2 ratio. Each column represents an independent experiment.

**Supplementary figure 3.**
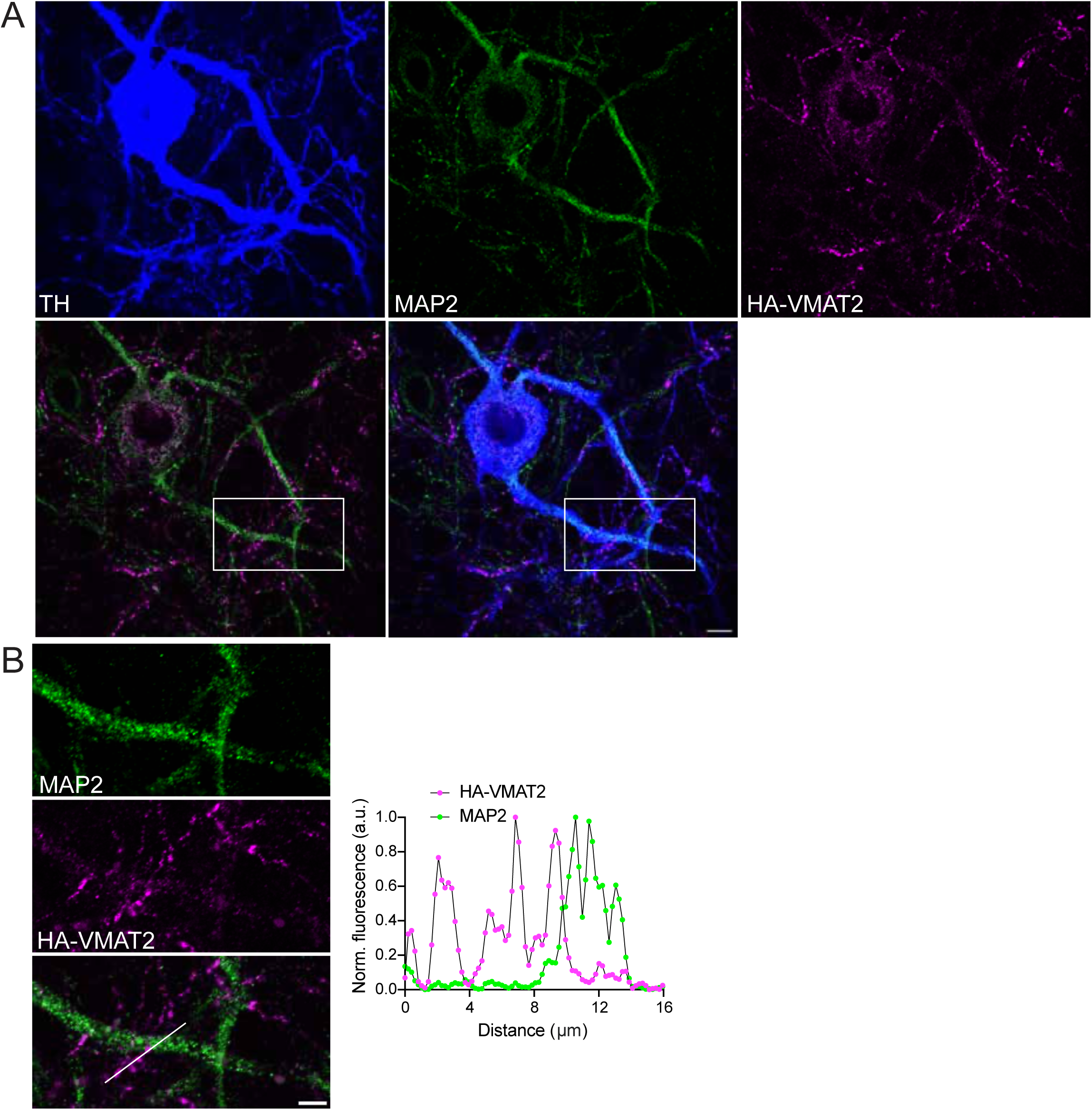
HA-VMAT2 varicosities represent axonal vesicle clusters. A. Confocal image of a dopamine neuron in a HA-VMAT2 KI midbrain culture stained for TH, somatodendritic protein MAP2 and HA-VMAT2. Scale bar = 10 µm. B. Magnified view of the area highlighted by white box in A. Scale bar = 5 µm. Graph shows normalized fluorescence profiles of HA-VMAT2 and MAP2 along the line ROI shown in the merged image.

**Supplementary figure 4.**
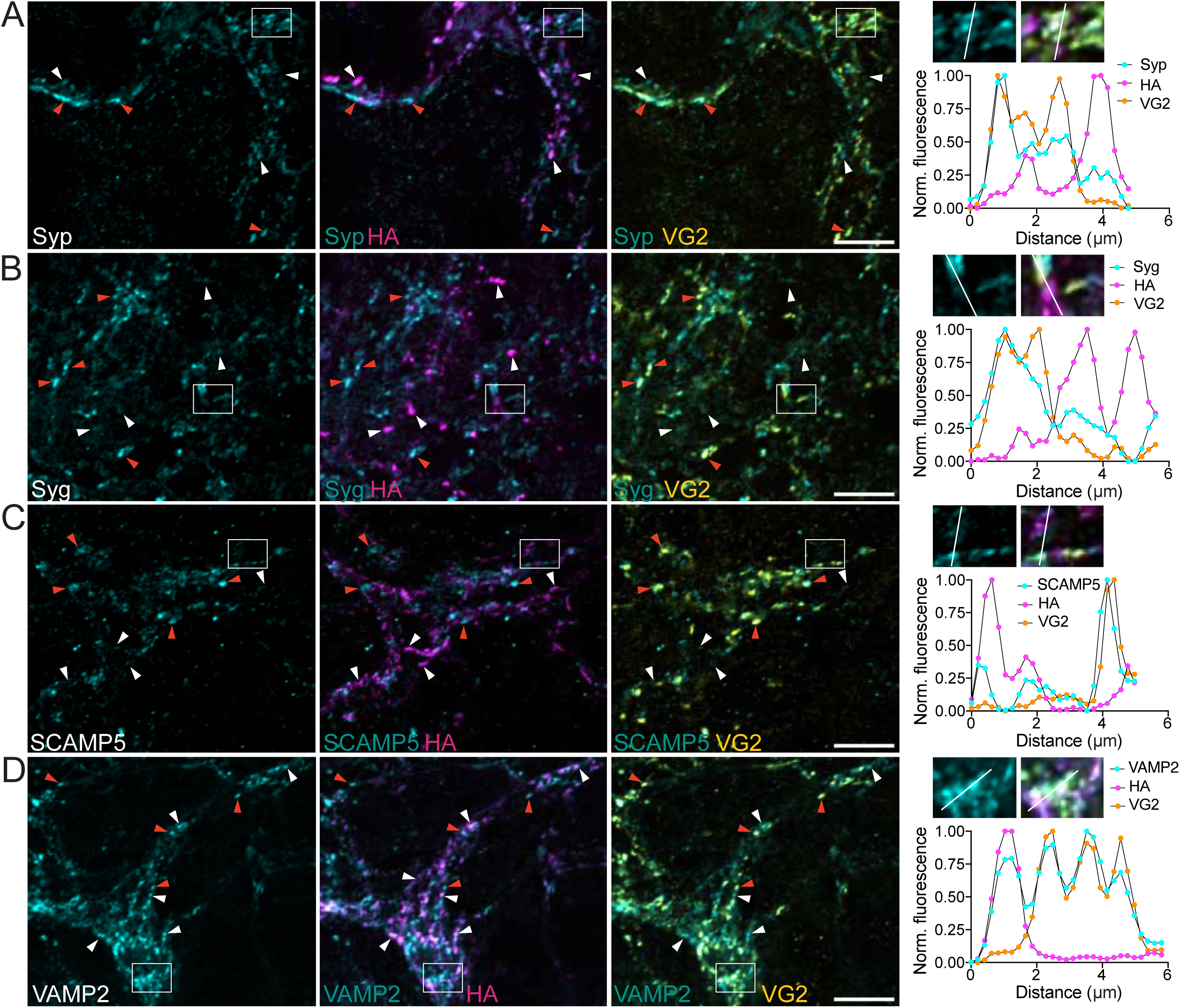
Presynaptic composition of dopamine and glutamate release sites in primary midbrain cultures. A-D. Confocal images of midbrain cultures from HA-VMAT2 KI mice stained for HA-VMAT2, VGLUT2 and synaptophysin (A), synaptogyrin1 (B), SCAMP5 (C) or VAMP2 (D). White arrowheads point to HA-VMAT2 and orange arrowheads to VGLUT2 varicosities. Scale bars = 10 µm in all images. Insets show magnification of the area indicated by the white boxes in larger images and a line ROI with corresponding fluorescence profiles in three channels shown below.

**Supplementary figure 5.**
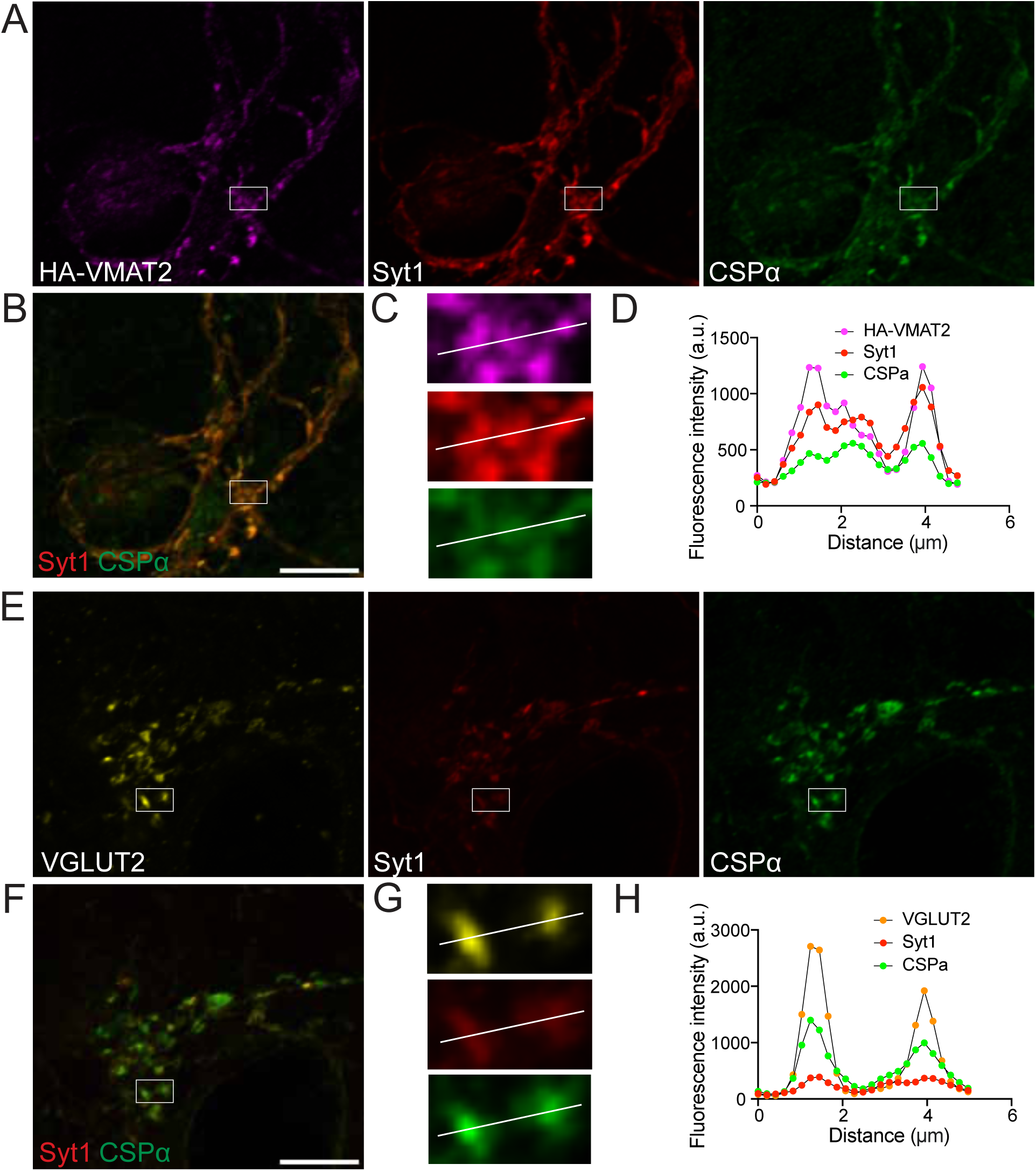
VMAT2 and VGLUT2 varicosities differ in syt1/CSPα ratio. A-D. A. Confocal image of a midbrain culture from HA-VMAT2 KI mice stained for HA-VMAT2, synaptotagmin1 and CSPα. B. Merge of synaptotagmin1 and CSPα channels. Scale bar = 10 µm. C. Magnified view of the area indicated by white box in A and B. D. Fluorescence profiles of HA-VMAT2, synaptotagmin1 and CSPα along the line ROI shown in C. E-H. E. Confocal image of a midbrain culture from HA-VMAT2 KI mice stained for VGLUT2, synaptotagmin1 and CSPα. F. Merge of synaptotagmin1 and CSPα channels. Scale bar = 10 µm. G. Magnified view of the area indicated by white box in E and F. H. Fluorescence profiles of VGLUT2, synaptotagmin1 and CSPα along the line ROI shown in G.

**Supplementary figure 6.**
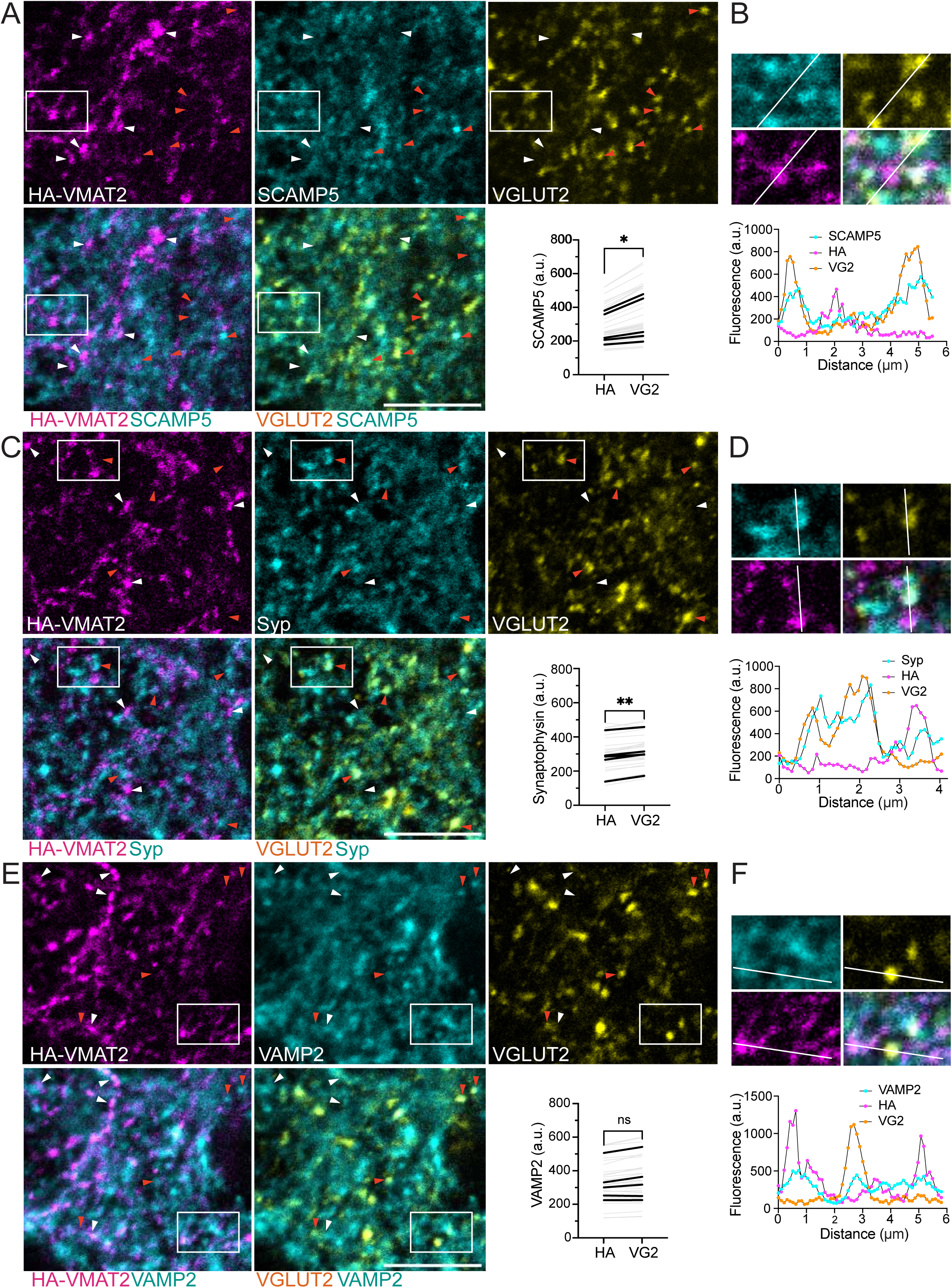
Protein composition of VMAT2 and VGLUT2 varicosities in the striatum. **A-B.** A. Confocal image of a single plane in a striatal slice from a HA-VMAT2 KI mouse stained for HA-VMAT2, SCAMP5 and VGLUT2 together with quantification of mean SCAMP5 intensity in 3D surfaces generated based on HA-VMAT2 or VGLUT2 signal. Gray lines show mean values of varicosities per image, black lines indicate the mean values per mouse. * = 0.0378 by paired t-test on mouse averages. Scale bar = 10 µm. White arrowheads point to HA-VMAT2 and orange arrowheads to a VGLUT2 varicosities. B. Magnified view of the area highlighted by white box in A and plot profile of fluorescence intensities in three channels along the line ROI. **C-D.** C. Confocal image of a single plane in a striatal slice from a HA-VMAT2 KI mouse stained for HA-VMAT2, synaptophysin and VGLUT2 together with quantification of mean synaptophysin intensity in 3D surfaces generated based on HA-VMAT2 or VGLUT2 signal. Gray lines show mean values of varicosities per image, black lines indicate the mean values per mouse. ** = 0.0043 by paired t-test on mouse averages. Scale bar = 10 µm. White arrowheads point to HA-VMAT2 and orange arrowheads to a VGLUT2 varicosities. D. Magnified view of the area highlighted by white box in C and plot profile of fluorescence intensities in three channels along the line ROI. **E-F.** E. Confocal image of a single plane in a striatal slice from a HA-VMAT2 KI mouse stained for HA-VMAT2, VAMP2 and VGLUT2 together with quantification of mean VAMP2 intensity in 3D surfaces generated based on HA-VMAT2 or VGLUT2 signal. Gray lines show mean values of varicosities per image, black lines indicate the mean values per mouse. P = 0.1129 by paired t-test on mouse averages. Scale bar = 10 µm. White arrowheads point to HA-VMAT2 and orange arrowheads to a VGLUT2 varicosities. F. Magnified view of the area highlighted by white box in E and plot profile of fluorescence intensities in three channels along the line ROI.

## METHODS

### Mice

Both male and female mice were used for all experiments. All mice were housed in group housing, received unlimited food and water and were exposed to 12h/12h light/dark cycle. HA-VMAT2 KI transgenic mice were used for proteomics, primary dopamine neuron cultures and tissue staining. C57BL6J mice were used for primary hippocampal cultures. All experiments with animals were performed according to the National Institutes of Health Guide for Care and Use of Laboratory Animals and were approved by the Cedars-Sinai Medical Center Institutional Animal Care and Use Committee.

### Generation of HA-VMAT2 knock-in mice

To generate the HA-VMAT2 knock-in mouse model, we used the CRISPR/Cas9 genome editing system. A guide RNA was designed to target the N-terminal tail of mouse VMAT2 (target sequence 5’-CAGCAGCACCAGATCGCTCA-3’). An ssDNA donor template was constructed to contain the sequence of the hemagglutinin tag followed by a linker consisting of two glycines and a modified PAM sequence. The donor sequence was flanked by 60 bp homology arms on both sides. Cas9 together with the gRNA and ssDNA donor were co-injected into C57BL/6 mouse embryos at a single cell stage at the University of California, San Francisco Transgenic Gene Targeting Core.

### Molecular biology

SCAMP5-specific small hairpin RNA (shRNA) was designed with the aid of the InvivoGen siRNA Wizard Online Tool to target the mouse SCAMP5 cDNA sequence. SCAMP5 knock-down (TGCTCTTAGCATGGTTCATAA) and a scrambled control sequences were inserted into the pLKO.1 lentiviral vector^99^ obtained from Addgene using AgeI and EcoRI restriction sites. An shRNA-resistant SCAMP5 rescue sequence was designed with the help of the Java Codon Optimization Tool, effectively creating silent mutations in the target region sequence which would be unrecognizable by the shRNA but would still code for SCAMP5. A custom lentiviral vector was ordered through Vector Builder and includes the human synapsin promoter, SCAMP5 rescue sequence, and mCherry to mark infected cells.

### Lentivirus production

Low-passage LentiX HEK cells (Takara, cat. # 632180) were seeded onto 6-well plates and transfected the following day with 1.5 μg of third-generation lentiviral vector FUGW encoding the gene of interest together with 0.5 μg each of packaging plasmids pRRE, pVSVG and pRSV-Rev using the FuGene HD transfection reagent (Promega, cat. # E2311) according to manufacturer’s instructions. Next morning, culture medium was changed into fresh Neurobasal. Viral supernatant was collected 47 hours after transfection and concentrated 10x using Lenti-X Concentrator (Takara, cat. # 631231). Aliquots were stored at −80C until use.

### Primary neuron culture

Hippocampal neurons: Hippocampi were dissected from P0 mice of both sexes in Hank’s Balanced Salt Solution (HBSS) containing 10 mM HEPES and 20 mM glucose, dissociated in 0.25% trypsin, washed three times with Minimal Essential Medium containing 1x B27 (GIBCO, 17504-044), 2 mM glutamax (GIBCO, 35050-061), 5% FBS (HyClone, defined), 21 mM glucose (Sigma, G8769) and 1x penicillin/streptomycin (Corning, 30-002-CL), triturated and plated on poly-L-lysine coated coverslips at 350 cells/mm2 in the above Minimal Essential Media. After one day *in vitro* (DIV1), 3/4 of the medium was changed to Neurobasal (GIBCO, 21103-049) with B27, glutamax and penicillin/streptomycin. Cells were infected with lentiviruses encoding VGLUT2 - pHluorin, VMAT2-pHluorin, and/or shRNA SCAMP5 KD, scramble or rescue on DIV 4-5. 4 μM cytosine arabinoside was added on DIV7 to inhibit glial proliferation.

Dopamine neurons: Ventral midbrain regions containing substantia nigra and ventral tegmental area were dissected from neonatal HA-VMAT2 KI mice of both sexes. Tissue pieces were dissected in Hank’s Balanced Salt Solution (HBSS) containing 10 mM HEPES and 20 mM glucose, dissociated in papain (Worthington) for 15 minutes, washed with media containing 60% Neurobasal-A, 30% Basal Media Eagle, 10% fetal bovine serum (HyClone, defined), 1x B27, 2 mM GlutaMAX, 10 ng/mL glial cell line-derived neurotrophic factor (Millipore, MA, USA) and 1x penicillin/streptomycin, triturated and plated in the above media at 1500 cells/mm^2^.

### Synaptic Vesicle Preparation

Whole brains or striatal tissue from 10-15 HA-VMAT2 KI mice was pooled per replicate. Tissue was homogenized by 9 strokes at 900 rpm in cold homogenization buffer (0.3M sucrose, 1mM Mg-EGTA, 10 mM HEPES-Tris, pH 7.4) with cOmplete protease inhibitors (Roche, cat. # 11836170001) using a glass teflon homogenizer. The homogenate (H) was centrifuged at 871 g for 10 min at 4 °C in an SS34 rotor (Sorvall). Pellet was discarded and supernatant (S1) was sedimented at 12,000 g for 15 min at 4 °C in an SS34 rotor. Both pellet P2 and supernatant S2 were collected and processed in parallel.

Pellet (P2) that is enriched in synaptosomes was resuspended in 10 mL homogenization buffer. Resuspended P2 was sedimented at 14,500 g for 15 min at 4 °C in an SS34 rotor, the supernatant (S2′) was removed and pooled with S2. P2′ was resuspended in 1.2 mL homogenization buffer and then subjected to osmotic lysis for 45 min in 13.5 ml cold water with 8 mM HEPES-Tris, pH 7.4. The lysed P2′ was homogenized using five strokes at 3,000 rpm and centrifuged at 32,500 g for 20 min. The lysed supernatant (LS1) was transferred into a new ultracentrifuge tube and centrifuged at 280,000 g for 2 h using a Beckman 70 Ti rotor. Supernatant was discarded and pellet (LP2) was resuspended in salt buffer (150 mM NaCl, 1 mM Mg-EGTA, 10 mM HEPES, pH 7.4) by repeated pipetting.

The S2/S2’ samples were homogenized by 5 strokes at 3000 rpm and sedimented 32,500 g for 20 min. The resulting supernatant S3 was transferred to a new ultracentrifuge tube and centrifuged at 280,000 g for 2 h using a Beckman 70 Ti rotor. Supernatant S4 was discarded and pellet P4 was collected and resuspended in salt buffer like LP2 described above.

To obtain pure SVs, LP2 and P4 fractions were further purified by velocity sedimentation through a continuous 5–25% glycerol gradient at 270,000 g for 1 h and 4 min in a Beckman SW 41 rotor. 0.5 mL fractions were collected from top to bottom. Fractions 7-12 were pooled and dialyzed over night at 4C in 150 mM NaCl, 1 mM Mg-EGTA, 10 mM HEPES, pH 7.4.

### Immunoisolation

Dialyzed synaptic vesicles were supplemented with 1% lysine and 1% glycine to reduce non-specific binding and cOmplete protease inhibitors (Roche, cat. # 11836170001). Vesicles were incubated on the rotator with anti-HA (Cell Signaling, Cat # 3724 for whole brain samples and Covance, AB_291262 for striatal samples), anti-VGLUT2 (Synaptic Systems, cat. # 95E11) antibodies for 2 hours at 4C. 5 μL of antibody was added per 2 mL of vesicle suspension.

In parallel, equal volumes of Dynabeads with protein A (Invitrogen 10001D) and protein G (Invitrogen 10003D) were washed three times in salt buffer with protease inhibitors and lysine/glycine. 50 μL of protein A/G beads was then added to 2 mL of synaptic vesicle/antibody mix and incubated on the rotator for 1 hour at 4C. Beads were washed three times in salt buffer and stored at −80C until processing for mass spectrometry.

### Mass Spectrometry

Peptide and protein identification and TMT quantitation: Peak lists were generated using the PAVA in-house software^100^. All generated peak lists were searched against the mouse subset of the SwissProt database (SwissProt.2019.07.31, 17,026 entries searched) using Protein Prospector^101^ with the following parameters: Enzyme specificity was set as Trypsin, and up to 2 missed cleavages per peptide were allowed. Carbamidomethylation of cysteine residues, and TMTPro16plex labeling of lysine residues and N-terminus of the protein were allowed as fixed modifications. N-acetylation of the N-terminus of the protein, loss of protein N-terminal methionine, pyroglutamate formation from of peptide N-terminal glutamines, and oxidation of methionine were allowed as variable modifications Mass tolerance was 5 ppm in MS and 30 ppm in MS/MS. The false discovery rate was estimated by searching the data using a concatenated database which contains the original SwissProt database, as well as a version of each original entry where the sequence has been randomized. A 1% FDR was permitted at the protein and peptide level.

For quantitation, only unique peptides were considered. Relative quantization of peptide abundance was performed by calculating the intensity of reporter ions corresponding to the different TMT labels present in MS/MS spectra. Intensities were determined by Protein Prospector. Median intensities of the reporter ions for all peptide spectral matches (PSMs) on each TMT channel were used to normalize individual (sample-specific) intensity values. In addition to classical SV proteins, we detected peptides associated with frequent contaminants of SV preparations (proteasomes, ribosomes). To improve detection of differences in SV proteins, contaminants associated with other organelles were excluded from analysis. For each PSM, relative abundances were calculated as ratios of signal in VGLUT2 versus matching VMAT2 samples. For relative amounts of total protein, peptide ratios were aggregated to the protein levels using median values of the log 2 ratios. Statistical significance was determined comparing the log2 ratios to a log2 reference value of 0 using a two-tailed t-test.

### Western blot

Proteins were separated by electrophoresis on 4-20% precast polyacrylamide gels (Bio-Rad) and transferred to nitrocellulose membranes. Membranes were blocked in 5% milk tris buffered saline with 0.1% Tween. Primary antibodies were incubated overnight at 4C and IRDye 680- or IRDye 800-coupled secondary antibodies (Rockland Immunochemicals) for 1 hour at room temperature.

### Immunohistochemistry

To obtain fixed brain, mice were perfused with cold PBS, followed by 4% PFA in PBS. Brains were postfixed overnight in 4% PFA and cryoprotected in PBS containing 30% sucrose. 35 μm sections were cut using a Leica CM3050 S cryostat and floating sections were stored in PBS with 0.02% Na-azide. Cultured cells were fixed for 15 minutes in 4% PFA in PBS and washed three times in PBS. Floating sections as well as cultured cells were blocked and permeabilized in PBS containing 5% normal goat serum and 0.2% Triton X-100. Primary and secondary antibodies were diluted in PBS containing 2.5% normal goat serum and 0.2% Triton X-100 in PBS for staining slices and in 2.5% normal goat serum in PBS for staining cultured cells. Primary antibodies were incubated overnight at 4°C for tissue sections and 2 hours at room temperature for cultured cells. Secondary antibodies were incubated with tissue sections for 2 hours and cultured cells for 1 hour in the dark at room temperature. Slices were mounted in Fluoromount-G (SouthernBiotech) or Vectashield (Vector Laboratories). Antibodies were used at the following dilutions:

SYT1 Ms 1:5000 Synaptic Systems 105011
TH Ms 1:1000 EMD Millipore MAB5280
VG2 Gp 1:1000 Synaptic Systems 135404
VG2 Ch 1:1000 Synaptic Systems 135416
SYT2 Gp 1:1000 Synaptic Systems 105225
HA Rt 1:500 Roche 11867423001
SYPH Ms 1:500 Synaptic Systems 101011
SYPH Gp 1:100 Synaptic Systems 101 308
VAMP2 Rb 1:500 Synaptic Systems 104202
CSP Rb 1:500 Synaptic Systems 154003
SLC10a4 Rb 1:200 Sigma HPA028835
SYG Rb 1:100 Synaptic Systems 103002
SCAMP5 Rb 1:50 Invitrogen PA5-61269
All secondary antibodies were used at 1/750 dilution.

### Imaging of fixed cultures and brain slices

Images of fixed cultured neurons and brain slices were acquired using the Nikon Eclipse Ti2-E Microscope equipped with AR or AX confocal, AR1-DUG2 scan head with gallium arsenide phosphide detector unit and a Plan Apo VC 60x DIC N2 oil objective. Lasers (405 nm, 488 nm, 561 nm and 640 nm) were used for excitation and ET525/50, ET593/46 and ET700/75 filters for emission. Colocalization within release sites of dopamine/glutamate coreleasing neurons was quantified on images acquired on a Nikon Eclipse Ti2-E microscope equipped with a AX confocal module and a NSPARC (Nikon Spatial Array Confocal) detector^60^. Lense alignment was re-calibrated every 45 minutes during imaging. Deconvolution was performed using Nikon’s built-in deblurring algorithm.

#### Tissue Section 3D Analysis

Presynaptic protein abundance in brain slices was quantified using Imaris software (Oxford Instruments). Z stacks were acquired with a z-step of 0.125 μm and 3D surfaces were generated based on HA-VMAT2 and VGLUT2 signal. Thresholds for surfaces were selected manually and kept consistent within-mouse for all analyses. For Syt-2 colocalization analysis, surfaces were generated manually and kept consistent within-mouse except for images of the lateral geniculate nucleus, where the morphology of VGLUT2 presynapses departed considerably from all other regions and unique thresholds were used. Mean signal for the protein of interest was measured in HA-VMAT2 versus VGLUT2 surfaces and obtained values were compared within image, and a paired t-test was applied for both slice- and mouse-level statistics.

### Image analysis in fixed cultures

Presynaptic protein abundance in midbrain cultures was quantified in imageJ. Regions of interest (ROI) were selected based on HA-VMAT2 or VGLUT2 signal or by the presence of both transporters. The fluorescence intensity of proteins of interest was measured in the three sets of ROIs. For colocalization analysis in dopamine/glutamate coreleasing neurons, images were processed using the Coloc2 plugin in imageJ.

### Live imaging

Hippocampal cultures between 16-19 DIV were mounted in a laminar flow perfusion and stimulation chamber (Warner Instruments RC-21BRFS) on an inverted Nikon Eclipse Ti fluorescence microscope, Hamamatsu ORCA-Flash4.0 camera, and Nikon Plan Apo *λ*D 60x oil objective. Fluorescence signals were collected under epifluorescence illumination using 465/95 nm excitation and 515/55 nm emission for pHluorin. Action potentials (APs) were evoked by passing 1 ms bipolar current pulses through platinum electrodes, to yield fields of 5–10 V/cm. Cells were continuously perfused with Tyrode’s buffer (119 mM NaCl, 25 mM HEPES, 2 mM CaCl_2_, 2 mM MgCl_2_, 2.5 mM KCl, and 30 mM glucose at pH 7.4) containing 10 μM 6-cyano-7-nitroquinoxaline-2,3-dione (CNQX) and 10 μM 3-(2-carboxypiperazin-4-yl)propyl-1-phosphonic acid (APV). Total cellular pHluorin content was revealed by applying Tyrode’s buffer with 50 mM NH_4_Cl, where NaCl was reduced to 69 mM. Imaging was done at 37°C. For analysis, synaptic boutons were selected manually. Background fluorescence was subtracted and fluorescence intensity in the synaptic boutons was normalized to pre-stimulation baseline and to unquenched (total) pHluorin fluorescence in NH_4_Cl.

### Statistics

Statistical analysis was performed using GraphPad Prism 10 software. The Shapiro-Wilk test was used to assess the normal distribution of data. For comparing two conditions, two-tailed paired or unpaired t-test (indicated in text and figure legends) was used if data was normally distributed and Mann-Whitney test was used if data that did not pass the test for normality. One-way ANOVA with Tukey’s post hoc test was used for comparing more than two samples. Data are presented as mean ± SEM and statistical significance as * P < 0.05, ** P < 0.01, *** P < 0.001, and **** P < 0.0001.

## ACKNOWLEDGEMENTS

This work was supported by grants from the Brain and Behavior Research Foundation (29845) and The Larry L. Hillblom Foundation (2022-A-004-SUP) as well as start-up funds from the Cedars-Sinai Medical Center to K.S. R.H.E. is supported by R01NS103938 and R01NS129803 from the National Institute of Neurological Disorders and Stroke. We are grateful for imaging support from V. Krishnan Ramanujan and Mingtian Che from the Cedars-Sinai Biobank and Research Pathology Resource. Mass spectrometry was performed at the Mass Spectrometry Resource at UCSF, which is supported by the Dr. Miriam and Sheldon G. Adelson Medical Research Foundation (J.A.O.-P. & A.L.B.).

## AUTHOR CONTRIBUTIONS

K.S. and R.H.E. designed research; H.A., A.J.D., H.X., J.A.O.-P., J.A., A.S., B.G., N.C. and K.S. performed experiments; H.A., A.J.D., H.X., J.A.O.-P., J.A., A.S., B.G. and K.S. analyzed data; A.L.B, R.H.E. and K.S. acquired funding; K.S. wrote the paper with feedback from R.H.E., H.A., A.J.D., H.X., J.A.O.-P. and A.S.

